# CRISPR-Cas9 cytidine and adenosine base editing of splice-sites mediates highly-efficient disruption of proteins in primary cells

**DOI:** 10.1101/2020.04.16.045336

**Authors:** Mitchell G. Kluesner, Walker S. Lahr, Cara-Lin Lonetree, Branden A. Smeester, Patricia N. Claudio-Vázquez, Samuel P. Pitzen, Madison J. Vignes, Samantha C. Lee, Samuel P. Bingea, Aneesha A. Andrews, Beau R. Webber, Branden S. Moriarity

## Abstract

Base editors allow for precise nucleotide editing without the need for genotoxic double-stranded breaks. Prior work has used base editors to knockout genes by introducing premature stop codons or by disrupting conserved splice-sites, but no direct comparison exists between these methods. Additionally, while base editor mediated disruption of splice sites has been used to shift the functional isoform pool, its utility for gene knockout requires further validation. To address these needs, we developed the program SpliceR (z.umn.edu/spliceR) to design cytidine-deaminase base editor (CBE) and adenosine-deaminase base editor (ABE) splice-site targeting guides. We compared the splice-site targeting and premature stop codon introduction in a knockout screen against the TCR-CD3 immune synapse in primary human T-cells. Our data suggests that 1) the CBE, BE4 is more reliable than the ABE, ABE7.10 for splice-site targeting knockout and 2) for both CBEs and ABEs, splice-donor targeting is the most reliable approach for base editing induced knockout.

## INTRODUCTION

Clustered Regularly Interspaced Short Palindromic Repeats (CRISPR) systems and their CRISPR associated proteins (Cas proteins) have allowed for an unprecedented ability to manipulate the genome (1–5). Key amongst the applications of these systems is their use in gene editing for targeted gene knock-out, knock-in, and modification^6^. These applications are of particular interest in the fields of cellular immunotherapies, where the multiplexed knock-out of genes involved in alloreactivity (TRAC, TRBC, B2M) as well as immunosuppressive genes such as *PDCD1, CTLA4, and TGFBR2*, in tandem with the knock-in of chimeric antigen receptors (CARs) specific to tumor antigens yields promise in the development of efficacious and safe therapies to recalcitrant malignancies^7–10^. In the most commonly used form of CRISPR, the Cas9 nuclease from *Streptococus pyogenes* (hereafter referred to as Cas9) is paired with a single-guide RNA (sgRNA) to induce a double stranded break (DSB) at a specific DNA site directed by the programmable complementarity of the 20nt sgRNA protospacer^5^. While CRISPR-Cas9 nucleases work exceptionally well for single gene editing, multiple concerns have emerged surrounding DSB induction, including large scale genomic rearrangements^11,12^ and altered pluripotency^13^, leading to the potential of oncogenesis in cell based therapies. Concerns arising from DSBs are exacerbated in a multiplex setting, where multiple genes are targeted simultaneously^7,14^. However, as more has been elucidated about the complex genetic circuitry involved in cancer immunosurveillance^15^, and the hope to generate ‘off the shelf’ CAR T cells^14^, it is of increased interest to modulate and edit genes in a multiplex setting^8^.

An alternative tool to edit genes without the need for DSBs are CRISPR-Cas9 base editors. Base editors are a class of gene editing enzymes that consist of a Cas9 nickase fused to a nucleotide deaminase domain^16–18^. In principle, base editors localize to a target region in the genome guided by the sgRNA protospacer. Once bound, the ssDNA R-loop is displaced by the binding of the Cas9 complex. Displacement allows the tethered deaminase domain access to the ssDNA R-loop, whereby cytidine deaminase base editors (CBEs, C-to-T) deaminate C-to-U, which base pairs like T, and adenosine deaminase base editors (ABEs, A-to-G) deaminate A-to-I, which base pairs like G. Nicking of the unedited strand by the core Cas9 nickase complex then stimulates DNA repair to use the nascent base as a template thus preserving the genomic edit during subsequent cycles of DNA replication.

Previously, our group established a platform for the multiplex engineering of human lymphocytes using CRISPR-Cas9 base editors. This approach achieves high-efficiency multiplexed gene knockout without multiple DSBs and their associated complications such as chromosomal translocations and hindered proliferation^14^. In that work we used CBEs to KO genes by mutating CAG, CGA, and TGG codons to introduce premature stop codons (pmSTOPs, previously termed iSTOP or CRISPR-STOP)^19,20^ and by mutating the conserved splice-site motifs to disrupt splicing of the fully functional transcript. To date, there has been substantial interest in using both CBEs and ABEs to modulate splicing which has predominantly focused on using targeted skipping of exons bearing pathogenic mutations (previously termed CRISPR-SKIP)^21,22^ and inducing functional alternative splicing patterns^23^. However, there is substantial evidence for the utility of the splice-site targeting methods for functional gene knockout as opposed to strictly modulating alternative splicing (Here, distinguished as BE-splice), which has been previously demonstrated with Cas9 nuclease^24,25^. Despite the array of reports demonstrating the various ways base editors can be used to knockout genes and modulate splicing, there are two main gaps in the field, namely 1) a comprehensive tool for designing both CBE and ABE sgRNAs to target both splice donor (SD) and splice acceptor (SA) sites, and 2) a head-to-head comparison of the methods of base editing mediated gene knockout encompassed across, ABEs, CBEs, BE-splice methods, and pmSTOP induction.

Here we present an easy-to-use webtool SpliceR (z.umn.edu/splicer), for the design of base editing sgRNAs to target splice-sites of any ensembl annotated metazoan genome using an ensembl transcript ID. We demonstrate the robustness of our approach in a focused sgRNA screen targeting the various proteins that make up the heterooctameric T cell receptor-CD3 (TCR-CD3) complex and MHC Class I immune synapse. We found that for gene editing and protein knockout 1) CBEs were more reliable than ABEs, 2) among both CBEs and ABEs targeting splice donors tended to produce more robust knockouts than targeting splice acceptors, and 3) targeting splice donor sites produced more robust knockouts than pmSTOPs. Ultimately, here we describe a highly-efficient method and program for the design of sgRNAs for gene knockout via base editing.

## RESULTS

In previous work, we and others have established the ability to target splice sites with base editing for both modulating splicing patterns and disrupting proteins^14,21,23^. While programs exist for designing splice acceptor targeting guides with a limited number of PAMs, there is not a comprehensive program for designing guides that target both splice donors and splice acceptors sites with CBEs or ABEs (Fig. 1a, 1b), that is compatible with any PAM identity, and that is compatible with all ensembl annotated metazoan species. To meet these needs, we developed the program SpliceR for the design of BE-splice sgRNAs (z.umn.edu/splicer, github repo). SpliceR employs an interactive user interface to specify the parameters for BE-splice sgRNA design, such as Ensembl transcript ID, PAM identity, base editor type, and species of interest (Supplementary Fig. 1a). SpliceR communicates directly with ensembl.org to query genetic information for sgRNA design, and then performs sgRNA matching and scoring.

**Figure 1.**
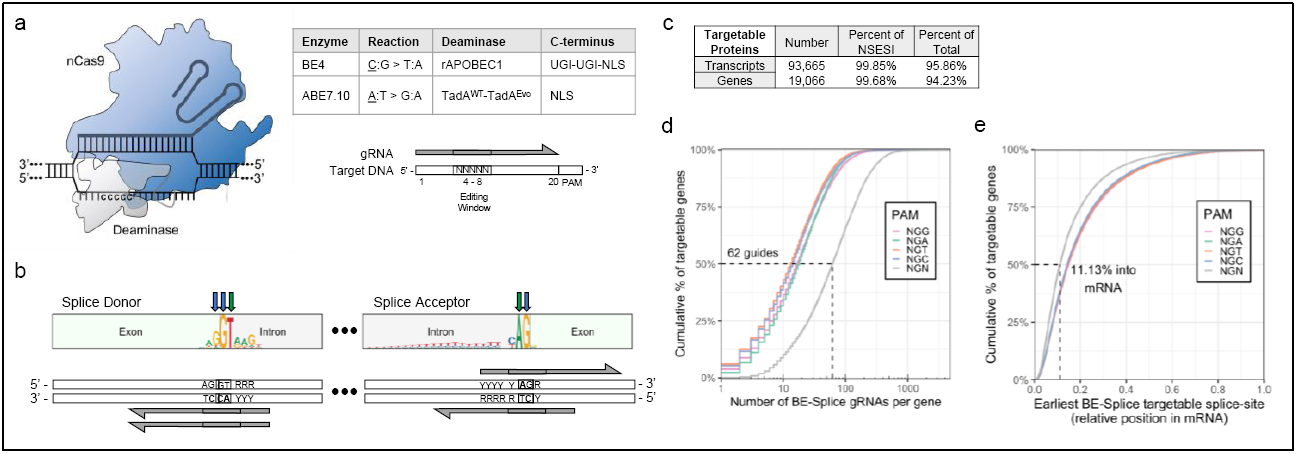
Overview of the BE-splice approach. (a) Diagram of base editor complex, table of the enzymes used in this study, and base editing sgRNAs with editing window. (b) Positioning of BE-splice sgRNAs within conserved splice donor and splice acceptors motif. Logo plots were generated from all human protein coding gene splice sites. Arrows indicate the base targeted by either CBEs (blue), or ABEs (green). (c) Breakdown of transcripts and genes targetable by BE-splice, showing the vast majority of non-single-exon-single-isoform (NSESI) genes are targetable by this approach (99.68%). (d) Distribution of BE-splice sgRNA density across each gene. 50% of genes have 62 or more sgRNAs mapping to them when accounting for all PAM identities and both CBE and ABE approaches. (e) Distribution of the position of the first sgRNA for each gene, with 50% having their first sgRNA 11.13% way through the mRNA or earlier.

To assess the practicality of the BE-splice approach, we used SpliceR to generate sgRNAs to every protein coding gene in the human genome. When restricting the analysis to genes that undergo splicing, i.e. non-single-exon-single-isoform (NSESI) genes, such that they are targetable by this approach, we found that 99.85% of transcripts and 99.68% of genes are expected to be targetable of the 94.23% of protein coding genes that are NSESI genes (Fig. 1c). Furthermore, 50% of NSESI genes are expected to have 62 sgRNAs or more, with the earliest sgRNA within the first 11.1% of the transcript (Fig. 1d, 1e). When broken down by targeting splice-donors or splice-acceptors, splice donors tend to have more sgRNAs (Supplementary Fig. 2), while also having the first sgRNA appear earlier in the transcript (Supplementary Fig. 3). These results suggested that the BE-splice approach is a robust method for targeting most genes, with targeting splice donors appearing more reliable given the greater availability of splice donor sgRNAs across CBEs and ABEs.

Despite the number of publications that have used base editors to disrupt or modulate genes by targeting splice sites with CBEs and ABEs, or introducing pmSTOPs with CBEs^14,19,20,22,23,26–28^, to our knowledge there is no comprehensive, direct comparison among these methods. Therefore, we sought to compare these methods directly and assess the predictions made from SpliceR in a mid-throughput screen. Given the interest of the TCR-CD3 and MHC class I immune synapse in the context of immunotherapies^29–33^, the presence of multiple NSESI genes that comprise these complexes (Fig. 2a), the necessity of every gene for surface expression (Fig. 2b, reviewed in Ref. 34^34^) and the ease of a functional readout with flow cytometry, we saw this complex as an ideal model to compare base editing mediated knockout approaches. To validate this model, we first designed Cas9 nuclease sgRNAs to each gene in the TCR-CD3 complex and the subunit of the MHC class I complex. Primary human T cells were then transfected with sgRNAs and Cas9 nuclease, and indel formation was measured via Sanger sequencing, while protein knockout was measured via flow cytometry (Supplementary Fig. 4). We found a high level of indel formation (81.7% ± 15.1%) and high corresponding protein knockout (73.4% ± 37.2%) across all genes in the complexes, including CD3ζ in the TCR-CD3 complex (Fig. 2c). These results establish that any one of the genes in the TCR-CD3 complex can be edited to induce a common phenotype of functional knockout.

**Figure 2.**
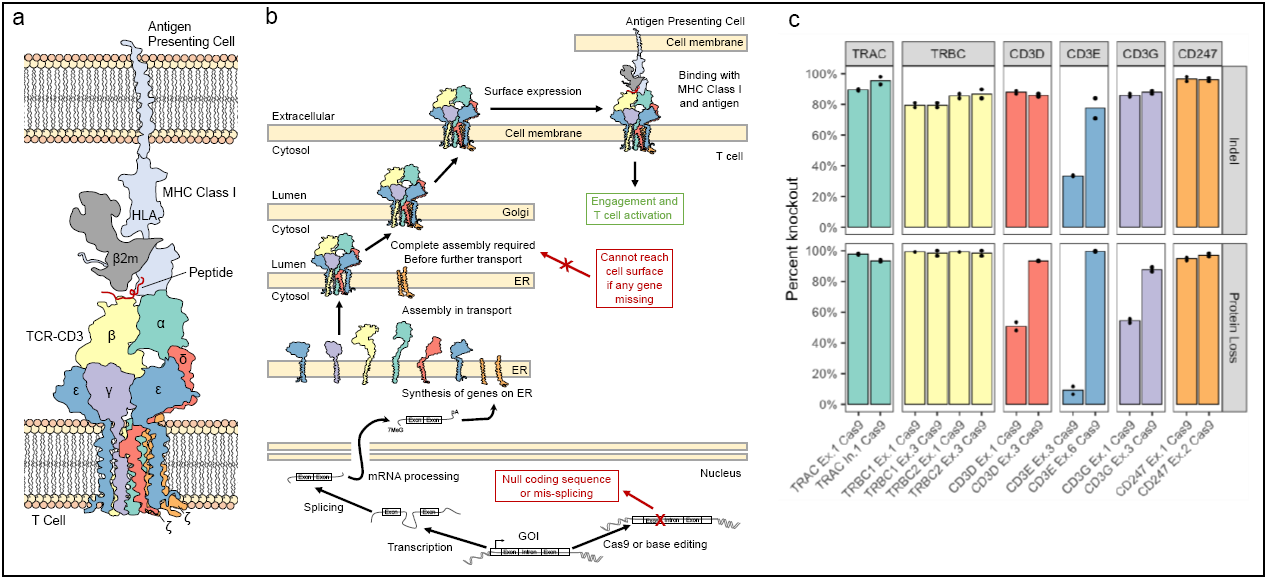
Conception and validation of the TCR-CD3-MHC Class I immune synapse as a screening model for functional knockout. (a) Diagram of the multimeric TCR-CD3 complex and MHC Class I immune synapse containing multiple NSESI genes, based on the solved structures (PDB 6JXR^43^, PDB 3T0E^47^; PDB 10GA^48^). (b) Diagram of the synthesis and localization of the TCR-CD3 complex and interaction with MHC Class I. All members of the CD3 complex are required before functional localization to the cell surface, where disruption of a single splice site within one gene member can prevent a surface expressed complex from forming. (c) Cas9 nuclease knockout of each individual member of TCR-CD3 complex validates the screening model. Two Cas9 nuclease sgRNAs were designed to exonic regions of each gene in the complex. All genes had at least one guide with >85% indel efficiency and loss in TCR-CD3 surface expression. N = 2 independent donors.

Next, we wanted to apply our model to directly compare 1) CBE vs. ABE knockout, 2) splice-donor vs. splice-acceptor knockout, and 3) BE-splice vs. pmSTOP knockout approaches. We used SpliceR to generate a screen of BE-splice sgRNAs targeting the TCR-CD3 complex, and the iSTOP database^19^ to design pmSTOP sgRNAs. Using the same workflow as with Cas9, primary human T cells were transfected with sgRNAs and either the cytidine base editor BE4, or the adenosine base editor ABE7.10, expressed via codon optimized mRNA^35^. Samples exhibited a wide range of gene editing at the target base across CBE and ABE approaches (M ± SD, 39.6% ± 31.6%, range 0%-100%), and a wide range of protein KO (M ± SD, 25.1% ± 31.9%, range 0%-97.0%) (Fig. 3a). Both editing of the target base and protein knockout was higher among CBE than ABE treated samples (t = 2.87, df = 76, p-value = 0.0053; t = 3.75, df = 76, p-value = 0.00034) (Fig. 3b, 3c).

**Figure 3.**
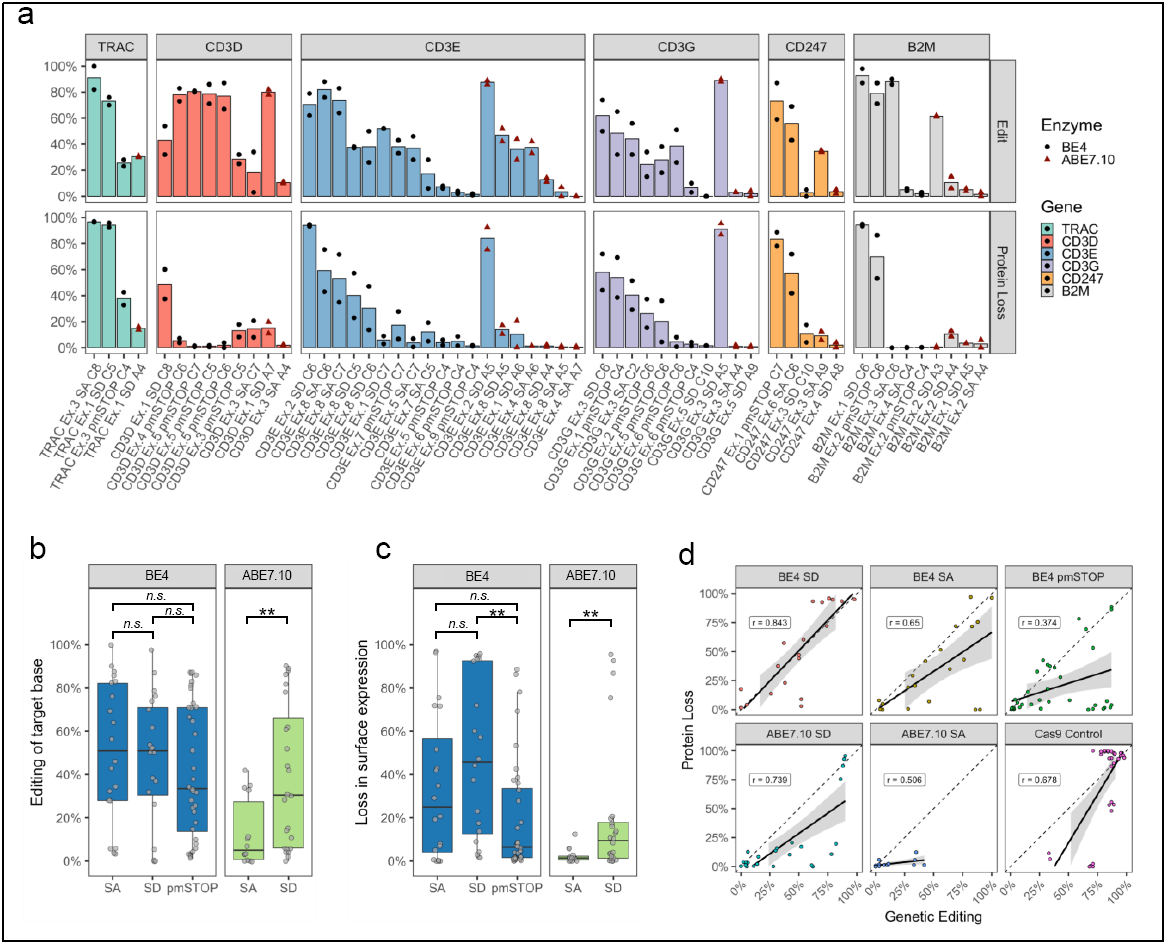
BE-splice sgRNAs mediate robust editing and disruption of TCR-CD3 MHC Class I immune synapse. (a) Editing efficiency (top) and surface protein loss (bottom) from each guide in the sgRNA screen. Results grouped by gene and enzyme used in descending order by protein loss. X-axis label indicates position of target base within sgRNA. TRBC1 and TRBC2 were omitted from the BE-splice screen due to the inability to design single BE-splice sgRNAs to target both paralogs simultaneously. (b) Base editing efficiencies grouped by enzyme and target motif analyzed with Student’s t-test or Welch’s t-test. (c) Protein loss efficiencies grouped by enzyme and target motif analyzed with Student’s t-test or Welch’s t-test. (d) Consistency of editing efficiency and protein loss across all approaches employed here. Relationship between protein loss and base editing efficiency is comparable to that observed in Cas9 control. N = 2 independent donors, performed on different days. *n.s.* P > 0.05, * P ≤ 0.05, ** P ≤ 0.01, *** P ≤ 0.001.

Within CBE treated samples, there was no significant difference in editing of the target base between splice donors, splice acceptors, and pmSTOPs (t = 0.416, df = 38, p-value = 0.680; t = -0.846, df = 54, p-value = 0.401; t = -1.27, df = 54, p-value = 0.204) (Fig. 3b). Furthermore, the rate of protein knockout was not significantly different between CBE splice donors and CBE splice acceptors (t = -1.27, df = 38, p-value = 0.213), however splice donors had significantly higher knockout efficiencies relative to pmSTOP sgRNAs (t = 3.33, df = 54, p-value = 0.00157), while splice acceptors were non-significantly higher than pmSTOP sgRNAs (t = 1.76, df = 54, p-value = 0.0841). Among CBEs, protein knockout was well correlated with editing of the splice donor and splice acceptor sites (r = 0.843, p-value = 3.11e-06; r = 0.65 p-value = 0.00192), yet poorly correlated with the introduction of premature stop codons (r = 0.374, p-value = 0.0247) (Fig. 3d). In contrast, ABE splice donor sgRNAs had significantly higher editing compared to ABE splice acceptors (t = -3.25, df = 34.9, p-value = 0.00257), which translated into the ABE splice donor protein knockout efficiency being significantly higher than the ABE splice acceptor protein knockout efficiency (t = -2.81, df = 23.9, p-value = 0.00972). Furthermore, protein knockout among ABE treated samples was well correlated with target base editing in splice donor (r = 0.739, p-value = 3.73e-05), but not in splice acceptor sgRNAs (r = 0.506, p-value = 0.0651). Among the well correlated BE-splice approaches; CBE splice donors, CBE splice acceptors, and ABE splice donor, the correlation was similar to that observed by Cas9 nuclease (r = 0.678, p-value = 2e-05) (Fig. 3d). Ultimately these results established that 1) CBE mediated knockout is more reliable than ABE mediated knockout, 2) splice donors are a more robust target across CBEs and ABEs, and 3) Among the genes in our screen, the BE-splice method produced more robust knockouts than pmSTOPs.

With these differences in mind, we wanted to investigate what may cause the disparities in baseline editing among the different approaches. Previous work has established that base editing efficiency is context dependent, with particular preference dictated by the nucleotide preceding the target base^36–38^. These works also sought to change the context dependencies of the preceding nucleotide by employing cytidine deaminase paralogs, orthologs, and engineered variants to change the context dependencies of base editors. Understanding the dinucleotide context dependencies of base editing would aid in the selection of BE-splice sgRNAs, however, to the best of our knowledge there does not exist a comprehensive analysis of the dinucleotide context dependencies by position of the commonly used base editors employed here, rAPOBEC1-BE4, and TadA_WT_-TadA_Evo_-ABE7.10.

To determine these dependencies, we performed an aggregated analysis for both BE4 and ABE7.10 with data across multiple cells types, genes and delivery methods from the literature^18,39–41^ and data generated by our group (6 papers, 102 guides, 447 edits in total). We chose to focus our analysis on editing efficiency as a function of the position in the protospacer and the identity of the preceding base. Aggregated analysis of BE4 across all nucleotide contexts produced a smooth distribution of editing activity centered about position 6 of the protospacer (Fig. 4a). Consistent with previous work, the editing window was dependent on the identity of the preceding nucleotide^17,26,37,42^, where T<u>C</u> dinucleotides exhibited the broadest editing window, while A<u>C</u> and C<u>C</u> exhibited smaller, comparable windows and G<u>C</u> exhibited a highly suppressed window with ≥ 20% activity only observed at positions 5-6 (Fig. 4a). When comparing T<u>C</u> to G<u>C</u> dinucleotides, the identity of the preceding nucleotide alone decreased the average editing efficiency by 3.2-fold from 28.7% to 6.9% (Supplementary Fig. 5a). In contrast, aggregated analysis of ABE7.10 yielded a narrower window with ≥ 20% editing activity between positions 4-8, along with thinner tails to the editing distribution (Fig. 4b). Interestingly, ABE7.10 exhibited similar preceding nucleotide context dependencies as BE4 with T<u>A</u> having the broadest and tallest window, followed by C<u>A</u>, A<u>A</u>, and then G<u>A</u> (Fig. 4b). When comparing T<u>A</u> to G<u>A</u> dinucleotides, the identity of the preceding nucleotide alone decreased the average editing efficiency by 2.9-fold from 24.5% to 6.3% (Supplementary Fig. 5b). Among both BE4 and ABE7.10, the post dinucleotide base did not appear to have as large of an effect on editing efficiency (Supplementary Fig. 5c-5f).

**Figure 4.**
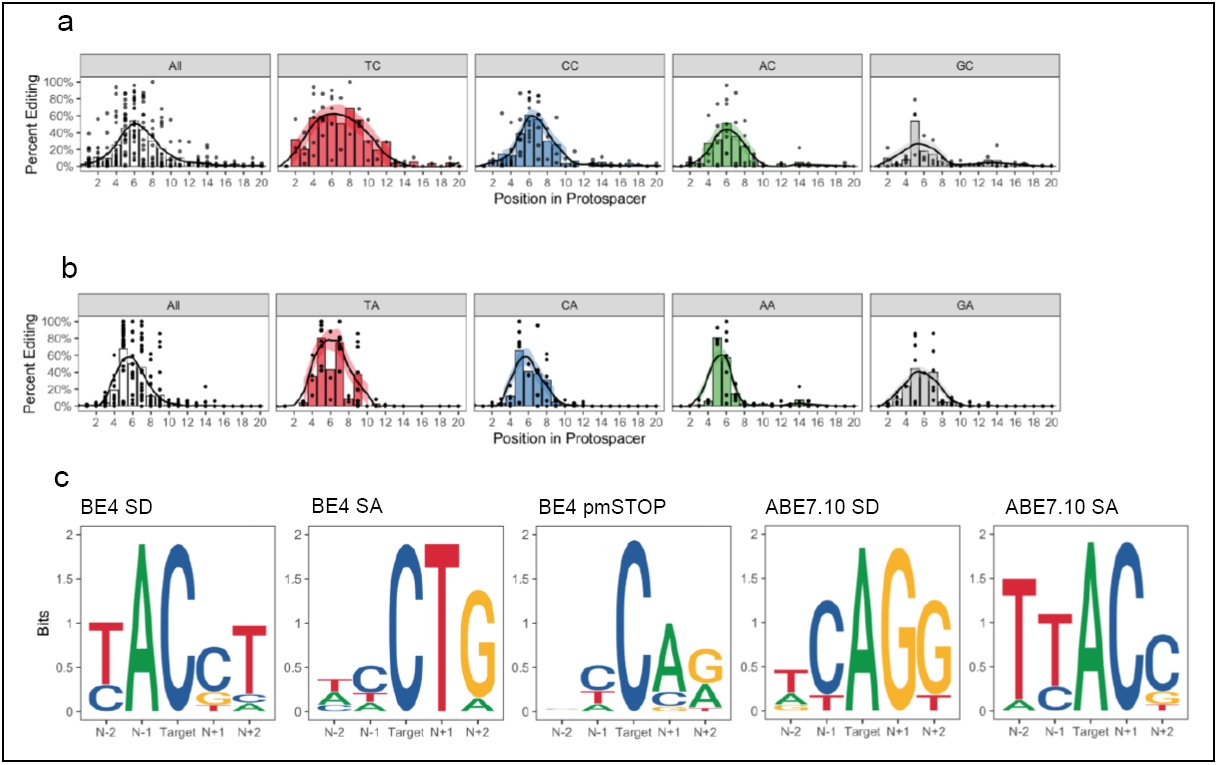
Context dependencies of base editors and BE-splice target motifs. (a) Preceding dinucleotide context dependencies of rAPOBEC1-BE4. Results normalized and aggregated across published results and our own work. Smoothed distributions generated using LOESS regression with span = 0.5. (b) Preceding dinucleotide context dependencies of TadA_WT_-TadA_Evo_-ABE7.10. Results normalized and aggregated across published results and our own work. Smoothed distributions generated using LOESS regression with span = 0.45. 6 papers, 102 guides, 447 edits. (c) Logo plots of pentanucleotide motif of each enzyme and target motif combination in this work. Plots are consistent with conserved splice site sequences.

Given the distinct differences in editing among dinucleotide contexts, we hypothesized that this could partially explain the differences in baseline editing efficiency of the BE-splice sgRNAs. To make this comparison we generated a consensus pentanucleotide motif for each approach we tested for base editing mediated protein knockout (Fig. 4c). Consistent with the expectation among CBE splice-site motifs, we observed an A<u>C</u>N motif for splice donors, N<u>C</u>T motif for splice acceptors, and a N<u>C</u>A motif for pmSTOPs. The lack of enhancing TC motifs, and inhibitory GC motifs across these approaches likely accounts for non-significant baseline editing efficiencies observed across all CBE treated samples. In contrast, ABE splice donors exhibited a preferred T<u>A</u>C motif, while the acceptors exhibited a non-disfavored C<u>A</u>G motif, which likely contributed to the significant difference in baseline editing among ABE treated samples (Fig. 3c).

Lastly, to investigate how the position of the sgRNA within the transcript affects the reliability of a knockout, we binned all base editing sgRNAs into first, second, middle, second-to-last, and last exons based on which exons they targeted (Supplementary Fig. 6). Strikingly, we found the highest degree of correlation between protein loss and base editing among guides that targeted middle exons (r = 0.915, p-value = 1.6e-20), and no significant correlation among guides that targeted the last exon (r = 0.359, p-value = 0.309), as might be expected (Fig. 5a). Next, we wanted to see how the error in protein loss, defined as the absolute value of the observed protein loss minus the editing of target base, varied as a function of the exons being targeted. We found that the error was minimized when targeting inner exons, and increased in guides targeting the second-to-last and last exons (Fig. 5b). Given the extracellular and transmembrane domains of the TCR-CD3 complex are essential to interchain interactions that assemble the complex^43^, allow for surface localization^34^, and signal transduction^44–46^, we wanted to study how the positioning of these sgRNAs affected the reliability of protein knockout. To do this, we mapped each sgRNA within the tertiary structure of the TCR-CD3 complex (PDB 6JXR^43^) and MHC Class I complex (PDB 3T0E^47^; PDB 10GA^48^), as determined by the solved structures (Supplementary Fig. 6-13). From these mappings, we classified each sgRNA as extracellular, intracellular, or transmembrane targeting. Consistent with the functional role of these regions, we found that guides targeting the transmembrane region of genes in the complex had the highest correlation of protein loss to editing (r = 0.966, p-value = 0.0017, albeit n = 6), followed by extracellular guides (r = 0.734, p-value = 1.18e-11), and intracellular guides (r = 0.435, p-value = 0.00314) (Fig. 5c). Ultimately, these results suggest that the greatest reliability of base editing mediated knockout is found targeting early, or inner exons within regions known to be functionally crucial.

**Figure 5.**
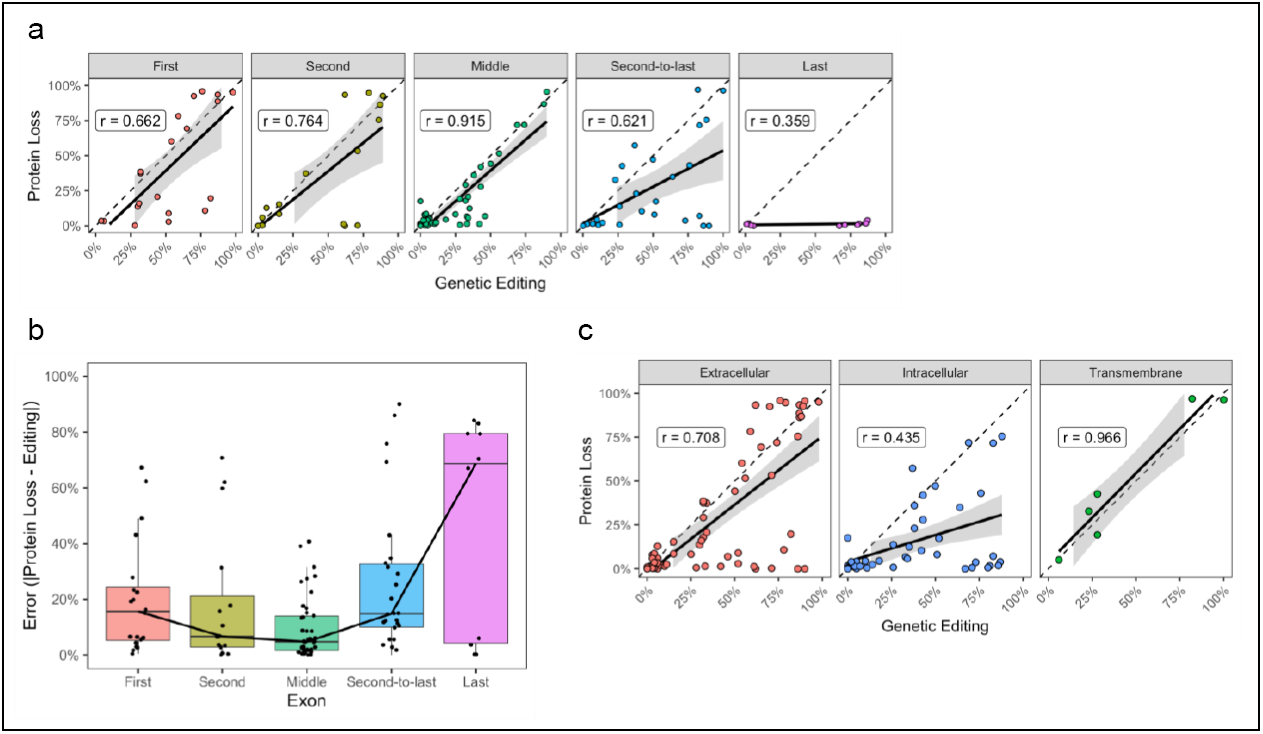
Consistency of editing efficiency and protein loss across mRNA and protein regions. (a) Scatter plots of protein loss and editing efficiency by exon group across all base editor approaches employed. The strongest relationship is observed among middle exons, while the weakest is observed in the last exon. (b) Error in protein loss as a function of editing efficiency across each exon group. The least error is observed among middle exons, while the greatest is observed in the last exon. (c) Scatter plots of the protein loss and editing efficiency grouped by where each BE sgRNA maps to the TCR-CD3 and MHC Class I structures. sgRNAs that map to transmembrane and extracellular regions exhibit the greatest consistency between protein loss and base editing efficiency.

## DISCUSSION

In this work we studied the use of CRISPR-Cas9 base editors for highly efficient gene disruption in primary human T cells using both cytidine and adenosine deaminase base editors. In this approach, the conserved splice donor (exon|GT-intron) and splice acceptor (intron-AG|exon) sites are edited via a transition mutation (C:G>T:A, or A:T>G:C), nullifying the splice-site and disrupting the gene at the transcriptional level^21,23^. To improve accessibility to the BE-splice approach, we developed the program SpliceR (z.umn.edu/splicer) as an online tool for the design of BE-splice sgRNAs. Analysis of the entire human genome showed that 95.86% of all protein coding gene transcripts, and 99.85% of all protein coding gene transcripts from genes that do not consist of a single-exon-single-isoform are targetable. We assessed these predictions, and compared the BE-splice approach to pmSTOP sgRNAs^19^ with a mid-throughput screen of sgRNAs targeting proteins involved in the TCR-CD3 MHC Class I immune synapse. From this screen we discovered three main trends, detailed below.

### Major findings

Firstly, CBE mediated knockout was more reliable than ABE mediated knockout. The higher rate of CBE knockout can be primarily attributed to the higher levels of baseline editing observed in CBE compared to ABE. The higher baseline editing of CBEs was consistent with the larger activity window of BE4 relative ABE7.10 observed in our aggregated analysis of these enzymes. Furthermore, across all dinucleotide contexts BE4 exhibited a smoother, more normally distributed window compared to ABE7.10, which may be the result of the processive activity of the CBE deaminase, *R. norvegicus* APOBEC1, while in contrast the ABE deaminase was evolved from *E. coli* TadA which acts endogenously on a single adenosine within the tRNA_Arg_ anticodon loop^49,50^. In general, we found current ABEs to be less reliable editors than CBEs, underpinning the need to develop more active ABEs by employing homologs, orthologs, and evolved variants as has been widely demonstrated among CBEs^17,26,37,42^. As such, during the final preparations of this manuscript two independent reports emerged describing the development of newer ABEs, termed ABE8s. These works employed either large-scale mutation libraries^51^ or phage assisted directed evolution ^52^ to produce ABEs with substantially higher on-target editing. Thus, employing these enzymes will likely reduce the disparity in baseline editing efficiency between CBEs and ABEs.

Secondly, across CBE and ABE, splice-donor sgRNAs produced more robust knockouts. In CBEs, splice donors and splice acceptors were not significantly different in editing efficiency, while in ABEs splice donors had a significantly higher editing efficiency. These trends were also mirrored in the rate of protein knockout with both enzymes, with the rate of protein loss being non-significantly higher in CBE splice donors relative to splice acceptors.

Among ABEs, this disparity could be attributed in part to the highly preferred T<u>A</u>C motif in splice donor guides, and the lesser preferred C<u>A</u>G motif in splice acceptor guides. The T<u>A</u>C preference motif is consistent with the preference motif of the parental enzyme *E. coli* TadA, and other adenosine deaminases^49,50,53^.

Additionally, previous work has demonstrated that targeting the splice donor sequence with an indel forming Cas9 nuclease causes robust protein and lncRNA knockout^24,25^. The reliability of disrupting the splice donor may be explained by the critical nature of the splice donor site in initiating splicing. During splicing, the pre-mRNA nucleotides contained in the splice donor site form a sequence-dependent RNA:RNA duplex with either the U1 or U12 snRNP. This Watson-Crick-Franklin interaction defines the exon boundary and initiates splicing^54,55^. Therefore, if this interaction is disrupted at the outset of splicing, then the ability to undergo the native splicing event may be principally inhibited.

Moreover, when considering the enhanced reliability of targeting splice donors from the splice acceptor perspective, previous work demonstrates that disrupting splice acceptors tends to favor exon skipping which may retain the reading frame, as opposed to a nonsense inducing outcome^21^. However, as demonstrated by the multiple instances of clinical splice site mutations (reviewed in Ref. 56^56^), it is important to note that splice acceptor mutations do not always result in a clean, single exon skipping outcome, as previous publications have suggested^21^. Rather, splice acceptor mutations are capable of inducing alternative splicing patterns via activation of cryptic splice sites, such as full intron retention, partial intron retention, and partial skipping of an exon^57–59^, all of which have the potential to knockout a gene by introducing a frameshift mutation, or removing a functionally critical region of the molecule.

Thirdly, among the genes in our screen, CBE-splice sgRNAs produced more frequent protein knockouts than pmSTOPs sgRNAs. At the genetic level, there was no significant difference in editing efficiency of BE-splice guides compared to pmSTOP guides, however, at the protein level pmSTOPs produced significantly less knockout compared to splice donors, and non-significantly less knockout compared to splice acceptors. These disparate results may be attributed to different levels of induction of nonsense mediated decay. Premature stop codons are known to incompletely induce a nonsense mediated decay^60^, or they can form relatively functional truncated variants^61^. In contrast, there is evidence that one-third of human genes have alternative isoforms that are actively suppressed by nonsense mediated decay^62^. This suggests that if a base edited splice site induces an alternative isoform that naturally occurs due to basal splicing errors, then these isoforms may be more readily subjected to nonsense mediated decay than newly introduced premature stop codons which have not been previously encountered by the cell.

Regardless of the mechanisms at play, the average knockout efficiencies of BE-splice and pmSTOP approaches could both be improved by increasing the baseline editing rates with more active CBEs such as EvoFERNY-BE4^37^, or hA3A-BE4^42^. Moreover, it is important to note that in our own view and application of using base editing to knockout genes, we see both the BE-splice and pmSTOP approaches as highly useful, complementary approaches in the screening of guides for protein knockout.

### Considerations for designing BE-splice sgRNAs and other BE-mediated knockout approaches

When designing BE-splice sgRNAs with SpliceR it is important to use the ensembl transcript table to choose whichever transcript or transcripts are most relevant to the biological phenomenon of interest. In instances where knockout of the main function of the gene is desired, we recommend targeting transcripts with merged Ensembl, Havana and Uniprot annotation (“gold labelled”). Furthermore, our results demonstrate that targeting all exons besides the last exon can produce high efficiency knockouts, where the innermost exons produce the most proportional protein knockout to genetic editing efficiency. We recommend initially screening 3-4 guides per gene with a greater than 50% predicted editing efficiency. If additional CBE sgRNAs are desired, we also recommend including a high predicted efficiency, inner exon pmSTOP sgRNAs to a known functional region, from the iSTOP database.

Furthermore, given situations where none of the aforementioned approaches are available, we also recommend trying to model known pathogenic variants such as those documented on ClinVar, Uniprot, or COSMIC for generating functional knockouts. However, as is the case with Cas9 nuclease editing, with any base editing mediated knockout approach we recommend functionally validating the effect of a base edit at the protein level, rather than assuming high editing efficiency corresponds to a functional knockout.

Lastly, much excitement has been generated by the prospect of prime editing^63^, which in principle allows for the precise editing and induction of mutations 80bp or smaller in size, including transversion, transition, and indel mutations. The ability to target a wide variety of mutations lends prime editing potential superiority over base editing in the correction of pathogenic mutations for gene therapy. This advantage is mainly derived from the ability of prime editing to precisely introduce desired mutations without undesired bystander edits to adjacent bases in the editing window as is seen with base editing. However, in the context of knocking out a gene by disrupting a conserved sequence, unintended bystander edits are of less concern when the goal is to nullify a splice site. Furthermore, despite the low-to-moderate level of indels observed in prime editing, this method still needs to be evaluated in a multiplex setting where compounding indels will likely lead to translocations^63^. The effect of these indels will also need to be addressed in stem cells, where double strand breaks have been associated with reduced potency^64,65^ and an increased risk of oncogenesis through inhibition of regulators the cell cycle and genomic stability^12,66^.

As more is understood about the deleterious consequences of Cas9 nuclease induced DSBs, it is of increasing interest to deploy gene editing methods that do not rely on DSBs. Here we show that cytidine deaminase base editors and adenosine deaminase base editors can allow for high efficiency knockout of proteins in primary T cells by disrupting conserved splice-sites. In a direct comparison of the major methods of base editing mediated knockout we validated with a sgRNA screen against the TCR-CD3 MHC Class I immune synapse. Our results demonstrate that 1) the CBE rAPOBEC1-BE4 is more reliable than ABE ABE7.10 for editing and knockout, and 2) across CBEs and ABEs, targeting splice donors is the most reliable method for base editing mediated knockout. Furthermore, in a composite analysis of data available from the literature as well as our own, we describe the dinucleotide contexts in detail of rAPOBEC1-BE4 and TadA_WT_-TadA_Evo_-ABE7.10. Collectively, these results inform the application of base editing for protein knockout, and in selecting optimal BE-splice guides generated by our program, SpliceR. Ultimately, we believe the BE-splice knockout approach for protein knockout is a widely applicable technique for genetic engineering, and cell based therapies, holding particular promise in the context of multiplex editing.

## METHODS

### T cell isolation

Peripheral blood mononuclear cells (PBMCs) were isolated from Trima Accel leukoreduction system (LRS, Memorial Blood Center) chambers using ammonium chloride-based red blood cell lysis followed by Ficoll-Paque gradient (GE Healthcare). PBMCs were further purified for CD3+ cells by immunomagnetic negative selection using the EasySep Human T-cell Isolation Kit (STEMCELL Technologies, Cambridge, MA). T cells were frozen at 10–20 × 10^6^ cells per 1 mL of Cryostor CS10 (STEMCELL Technologies, Cambridge, MA) and thawed into culture for knockout experiments.

### Cell culture

Primary human T cells were cultured in OpTmizer CTS T cell Expansion SFM (ThermoFisher, Waltham, MA) containing 2.5% CTS Immune Cell SR (ThermoFisher), L-Glutamine, Penicillin/Streptomycin, N-Acetyl-L-cysteine (10 mM, Sigma-Aldrich), IL-2 (300 IU/mL, PeproTech), IL-7 (5 ng/mL, PeproTech), and IL-15 (5 ng/mL, PeproTech) at 37 °C and 5% CO2. Prior to electroporation T cells were activated with Dynabeads Human T-Activator CD3/CD28 (ThermoFisher, Waltham, MA) at a 2:1 bead:cell ratio for 48 h. Following electroporation, T cells were maintained at 1 × 10^6^ cells/mL in a 24-well or 12-well plate.

### Transfection

Forty-eight hours post activation, Dynabeads were magnetically removed and cells were washed one time with PBS prior to resuspension in the appropriate electroporation buffer. T cells were electroporated with 1 µg of chemically modified sgRNA (Synthego, Menlo Park, CA) and 1.5 µg codon optimized BE4 mRNA (TriLink Biotechnologies, San Diego, CA) codon optimized ABE7.10 mRNA (TriLink Biotechnologies, San Diego, CA), or codon optimized *Sp*Cas9 mRNA (TriLink Biotechnologies, San Diego, CA). Cas9 nuclease sgRNAs were designed using the Synthego knockout guide design (https://design.synthego.com/#/), BE-splice sgRNAs were designed using SpliceR (z.umn.edu/splicer), and pmSTOP sgRNAs were designed using the iSTOP database (http://www.ciccialab-database.com/istop). T cells were electroporated with the 4D-nucleofector system (Lonza, Basel, Switzerland) using a P3 16-well Nucleocuvette kit, with 1 × 10^6^ T cells per 20 µL cuvette using the program EO-115. T cells were allowed to recover in electroporation cuvettes for 15 minutes before resuspension in antibiotic-free medium at 37 °C, 5% CO2 for 20 min following gene transfer, and were then cultured in complete CTS OpTmizer T cell Expansion SFM as described above.

### Flow cytometry

Seven days after electroporation a total of 5 × 10^6^ T cells were collected and stained in 100 uL with fluorophore-conjugated anti-human antibodies for CD3 (BD Biosciences #564001), beta-2-microglobulin (BioLegend #316306) and Fixable Viability Dye eFluor780 or LIVE/DEAD Fixable Aqua Dead Cell Stain (ThermoFisher #65-0865-14 1:500 dilution, #L34966 1:40 dilution) were used to assess cell viability. Events were acquired on LSR II or LSRFortessa flow cytometers using FACSDiva software, and data was analyzed using FlowJo v10 software. See Supplementary Fig. 4 for gating strategy. The percent of positive events were normalized, by dividing over the percent positivity seen in pulse alone controls. Protein loss was then calculated as [1 - % normalized sample positivity].

### Genetic analysis

Seven days after electroporation Genomic DNA was isolated from T cells by spin column-based purification (GeneJET, ThermoFisher). Editing efficiency was analyzed on the genomic level by PCR amplification of the targeted loci (Additional Data 7 for primers and guide protospacer), Sanger sequencing of the PCR amplicons (ACGT or Eurofins Genomics). Sanger sequencing traces of base edited samples were analyzed using EditR (z.umn.edu/editr)^67^, while indel efficiency was analyzed using the Synthego ICE (https://ice.synthego.com/#/)^68^.

### SpliceR development and whole genome guide prediction

SpliceR was written in the statistical programming language R (v. 3.6.1) using RStudio (v. 1.1.383). The SpliceR web app was developed using R shiny (https://shiny.rstudio.com/). All files are available online (https://github.com/MoriarityLab/SpliceR); dependencies.R is responsible for installing the necessary libraries to run the application; global.R defines the functions needed to interact with ensembl, and generate BE-splice sgRNAs. server.R and ui.R communicate with each other to handle the user inputs, generate sgRNAs, and return the output to the user interface. SpliceR relies on the various tidyverse (https://www.tidyverse.org/) and Bioconductor (https://www.bioconductor.org/) packages. For the whole genome guide prediction, the *generate_guides()* function from the SpliceR web app was modified to run as a command line executable. All human protein coding gene ensembl transcript IDs were pulled from the GENCODE database (https://www.gencodegenes.org/human/). The generate_guides() function was then run in parallel across all Ensembl transcript IDs using the Minnesota Supercomputing Institute (MSI).

### Aggregate analysis of CBE and ABE context

Papers employing the CBE rAPOBEC1-BE4, and the ABE TadA_WT_-TadA_Evo_-ABE7.10 were found using PubMed and Google scholar. Using a combination of main and supplementary figures the editing values for each targetable base (adeninosines for ABEs, and cytidines for CBEs) within the protospacer was manually recorded. Data generated from our own work was similarly entered. All manual data entries were double entered over two independent times to check for consistency of values. To control for variable electroporation and baseline editing efficiencies across different works, each editing value was normalized to the maximum editing efficiency observed in each cell type, for each work. To account for the same guide being used in different studies, normalized percent editing was averaged across each position of each unique guide. Data was then analyzed based on the pre-dinucleotide, and post-dinucleotide context of each base. R script for reproducible analysis is available in additional file 1, with aggregate analysis data from additional file 4 and additional file 5.

### Data analysis and visualization

All statistical analyses were performed in R using RStudio. The level of significance was set at α = 0.05. All statistical tests were first subjected to assumptions of homoscedasticity. For samples with equal variance, Student’s two-sample, unpaired two-tailed t-test was used, while for samples with unequal variance Welch’s two-sample, unpaired two-tailed t-test was used. Data were visualized in R studio employing various tidyverse (https://www.tidyverse.org/), Bioconductor (https://www.bioconductor.org/) packages, and ggseqlogo (https://github.com/omarwagih/ggseqlogo) packages. See R script in additional file 1 for reproducible analyses.

## DATA AVAILABILITY

Data and scripts for analysis and figure reproduction are available on GitHub. Source data are available from corresponding author upon request.

## ACKNOWLEDGEMENTS

We thank Dr. Matthew Johnson at the University of Minnesota for discussions surrounding flow cytometry. We thank Dr. Kevin Holden at Synthego for insight about chemically modified sgRNAs.

## Contributions

M.G.K., W.S.L., B.S.M., and B.R.W. designed experiments. M.G.K., W.S.L., C.L.L., B.A.S., P.N.C., M.J.V., S.C.L., S.P.B., and A.A.A. performed experiments. M.G.K., W.S.L., C.L.L., S.P.B., S.P.P., and S.C.L. analyzed the data. M.G.K. wrote the program. M.G.K., W.S.L., B.S.M., and B.R.W. wrote the manuscript. All authors approved and read the final manuscript.

## SUPPLEMENTARY INFORMATION

1. Supplementary figures
2. Script for figure reproduction of analyses and figures
3. Whole genome BE-splice sgRNA prediction data
4. Experimental data for each sample
5. CBE aggregate analysis data
6. ABE aggregate analysis data
7. SnapGene maps of genes with annotated BE-splice sgRNAs
8. Primer sequences for each sgRNA

