## Supplementary Figures for "CRISPR-Cas9 cytidine and adenosine base editing of splice-sites mediates highly-efficient disruption of proteins in primary cells"

Figures
Updated April 3rd, 2020

**AUTHORS**

Mitchell G. Kluesner[1,2,3,4]*, Walker S. Lahr[1,2,3,4],* Cara-Lin Lonetree[1,2,3,4], Branden A. Smeester[1,2,3,4], Patricia N. Claudio-Vázquez[1,2,3,4,5], Samuel P. Pitzen[5], Madison J. Vignes[6], Samantha C. Lee[6], Samuel P. Bingea[1,2,3,4], Aneesha A. Andrews[6], Beau R. Webber[1,2,3,4,†], Branden S. Moriarity[1,2,3,4,†]

*These authors contributed equally
†These authors contributed equally

**AFFILIATIONS**
[1]Department of Pediatrics, University of Minnesota, Minneapolis, MN, USA
[2]Masonic Cancer Center, University of Minnesota, Minneapolis, MN, USA
[3]Center for Genome Engineering, University of Minnesota, Minneapolis, MN, USA
[4]Stem Cell Institute, University of Minnesota, Minneapolis, MN, USA
[5]Department of Genetics, Cell Biology, and Development, University of Minnesota, Minneapolis, MN, USA
[6]College of Biological Sciences, University of Minnesota, Minneapolis, MN, USA

### Supplementary Figures

Figure S1

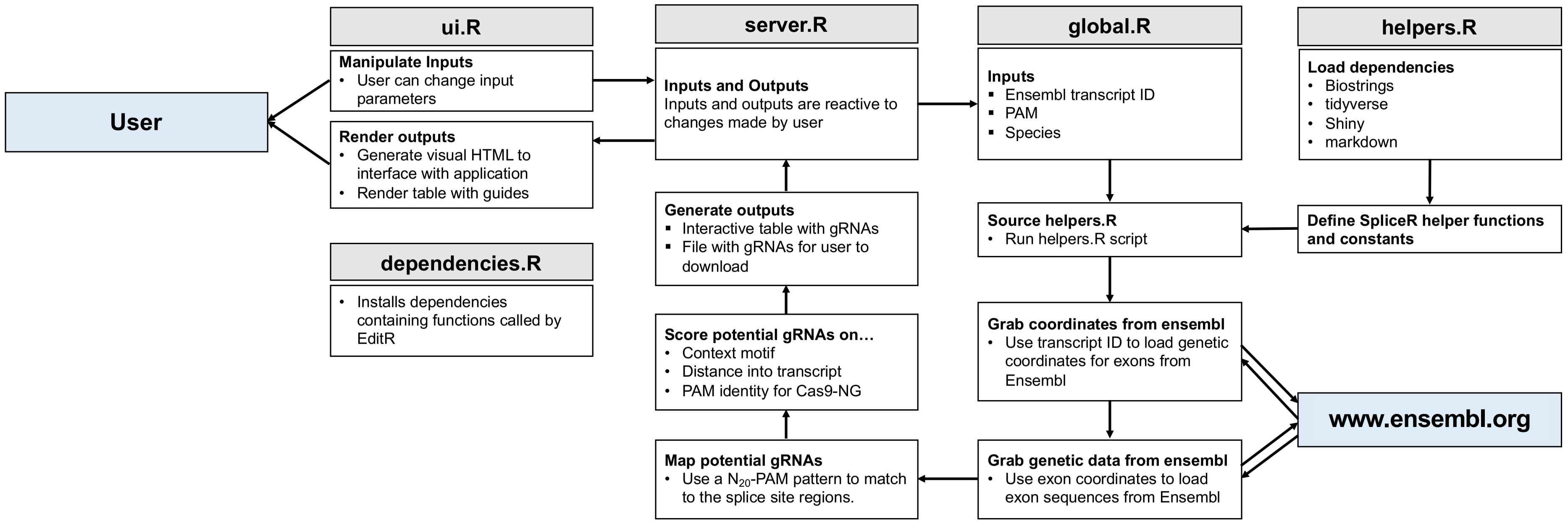


**Supplementary Figure 1**. Diagram of spliceR algorithm located at z.umn.edu/splicer. User provides an ensembl transcript ID, a preferred PAM, and the species of interest to target and SpliceR communicates directly with Ensembl to pull genetic information and map guides. sgRNAs are then scored by context motif, distance into transcript and PAM identity. Users can download predicted sgRNAs.

### Figure S2

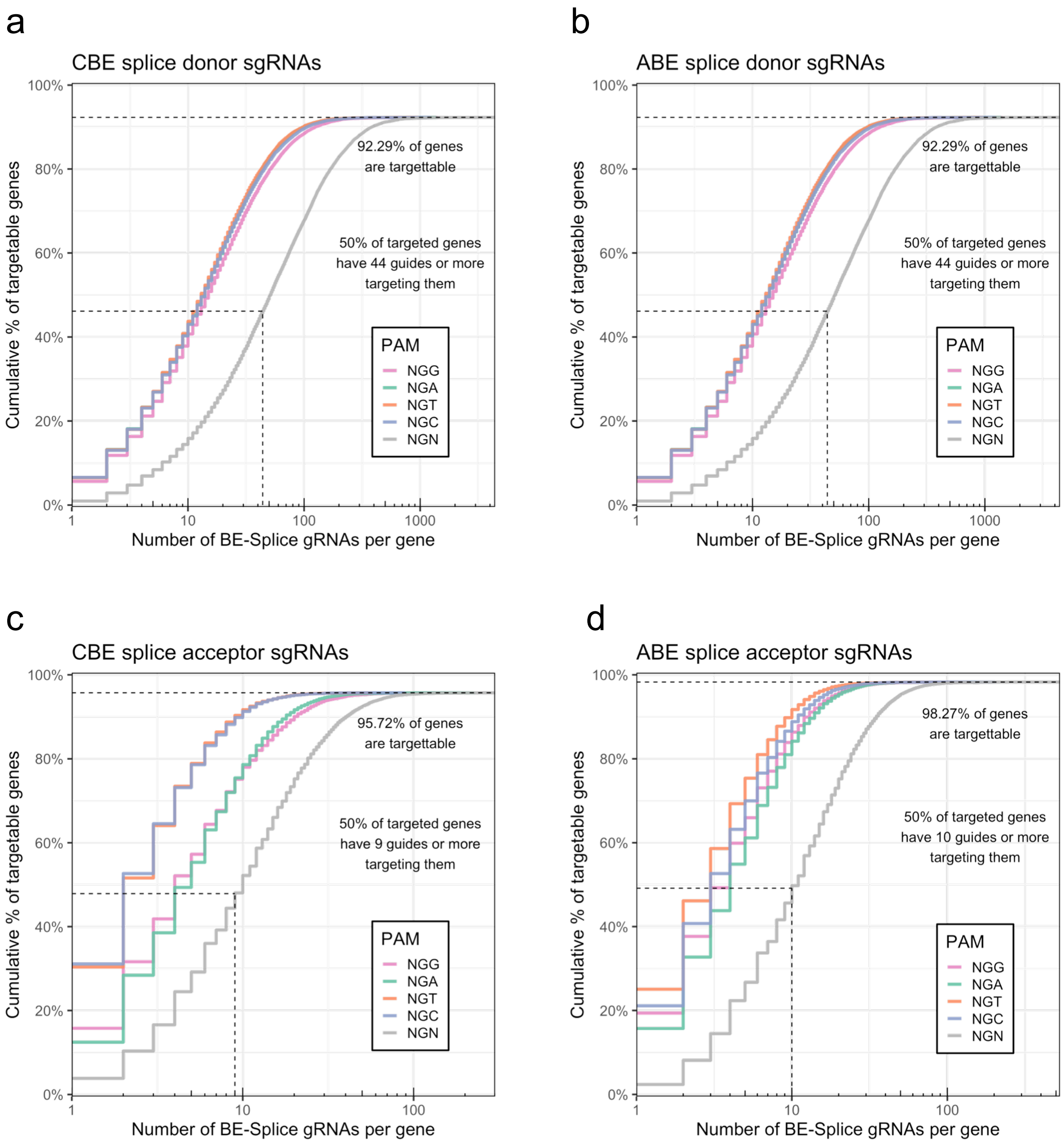


**Supplementary Figure 2.** Distribution of BE-splice sgRNA density across all genes by BE-splice approach. (a) CBE splice donors, (b) ABE splice donors, (c) CBE splice acceptors, (d) ABE splice acceptors. Note that CBE and ABE splice donors utilize the same sgRNAs, hence the same guide density.

### Figure S3

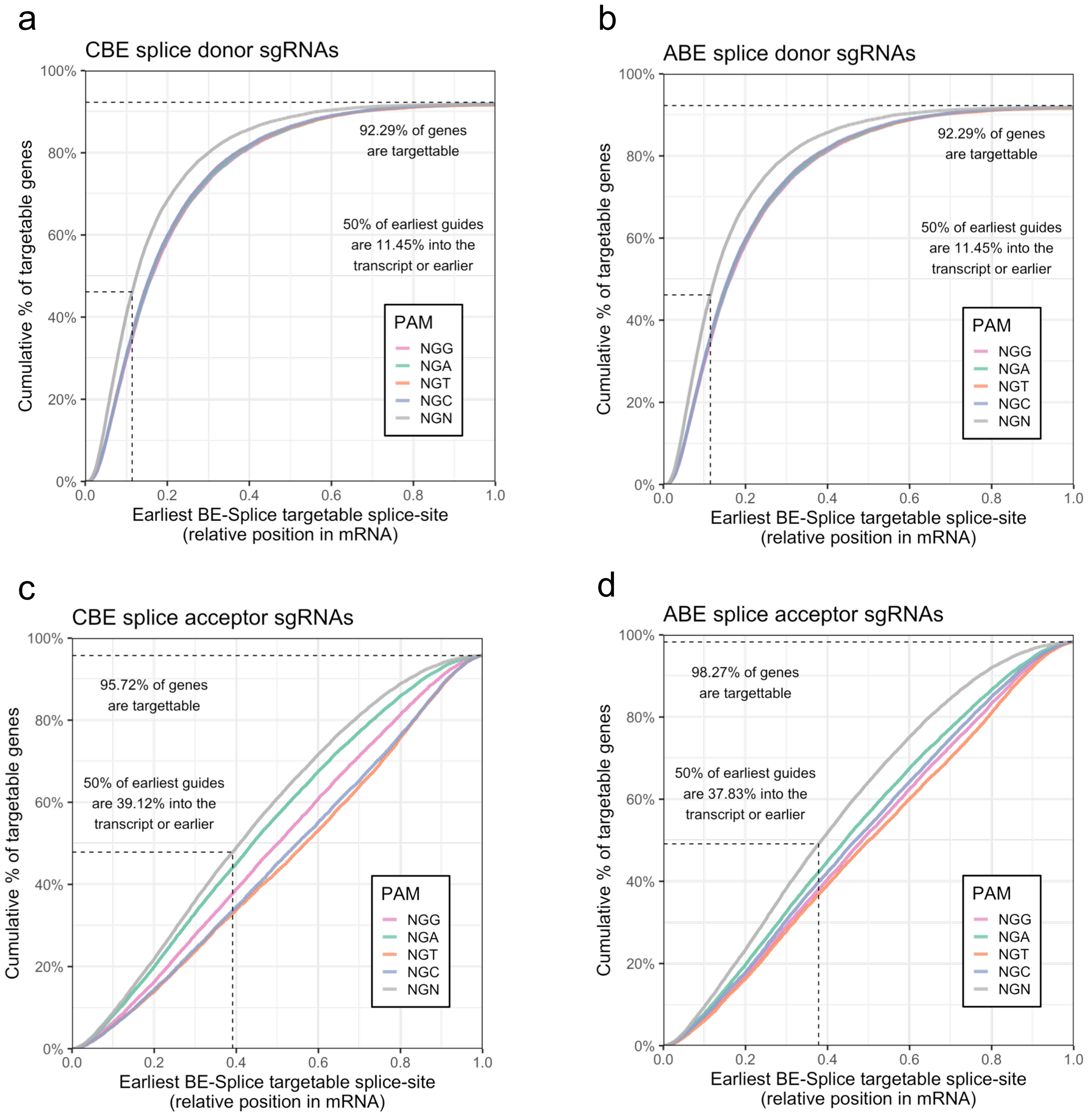


**Supplementary Figure 3.** Distribution of the position of the first sgRNA across all genes by BE-splice approach. (a) CBE splice donors, (b) ABE splice donors, (c) CBE splice acceptors, (d) ABE splice acceptors. Note that CBE and ABE splice donors utilize the same sgRNAs, hence the same guide density.

### Figure S4

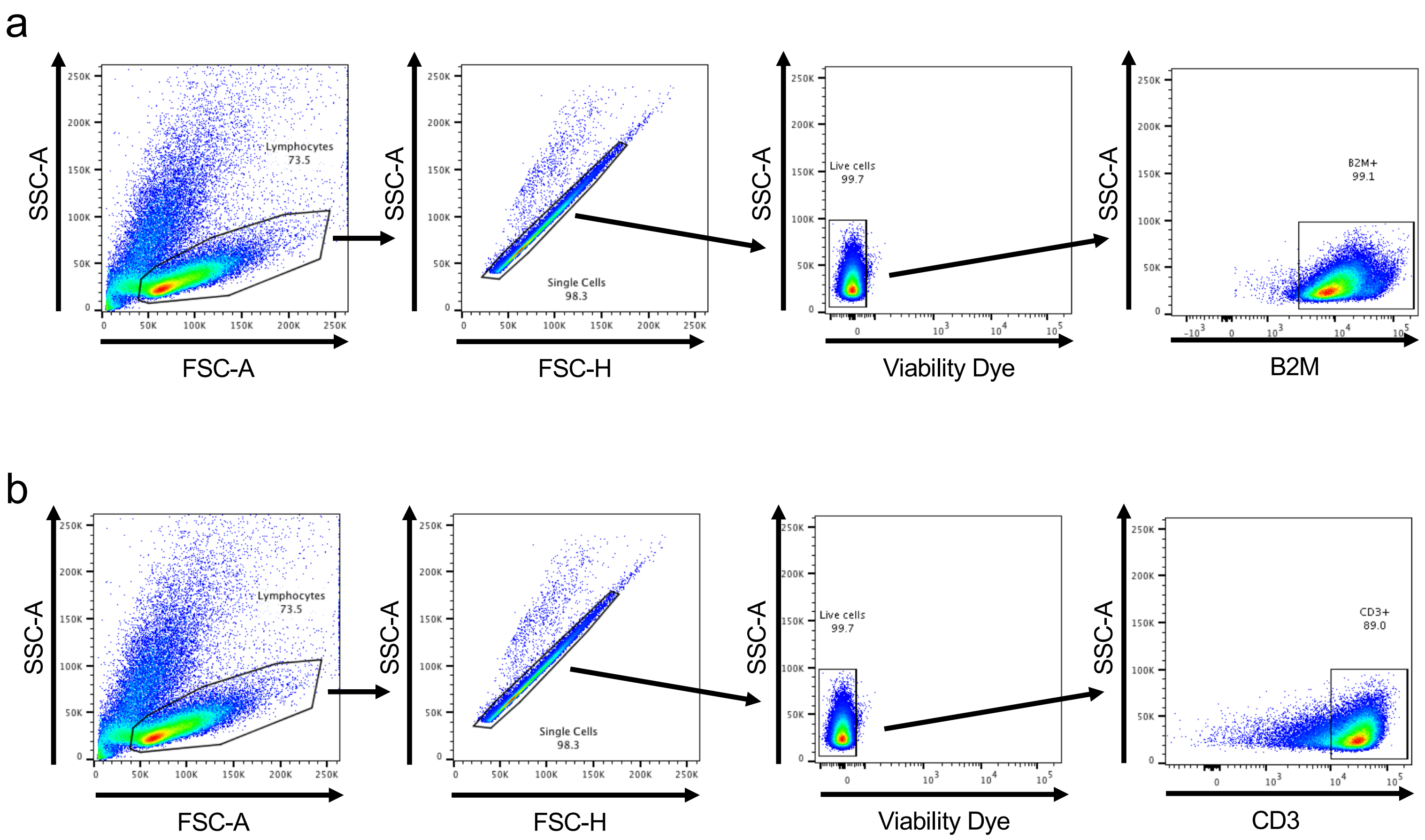


**Supplementary Figure 4.** Representative gating strategies for flow cytometry. (a) Gating tree for B2M$^+$ cells. (b) Gating tree for CD3$^+$ cells.

### Figure S5

a

| Dinucleotide | Average Editing | Std. Error | t value | Pr(>\|t\|) |
|---|---|---|---|---|
| All | 17.4% | 1.42% | 12.20 | 0.00e+00 |
| TC | 28.7% | 3.07% | 9.33 | 0.00e+00 |
| CC | 21.1% | 2.68% | 7.88 | 0.00e+00 |
| AC | 13.8% | 2.91% | 4.73 | 2.70e-06 |
| GC | 6.9% | 3.05% | 2.25 | 2.48e-02 |

b

| Dinucleotide | Average Editing | Std. Error | t value | Pr(>\|t\|) |
|---|---|---|---|---|
| All | 12.6% | 1.33% | 9.45 | 0.000000 |
| TA | 24.5% | 2.93% | 8.37 | 0.000000 |
| CA | 13.5% | 2.73% | 4.93 | 0.000001 |
| AA | 10.2% | 2.70% | 3.78 | 0.000170 |
| GA | 6.3% | 2.47% | 2.55 | 0.010800 |

c

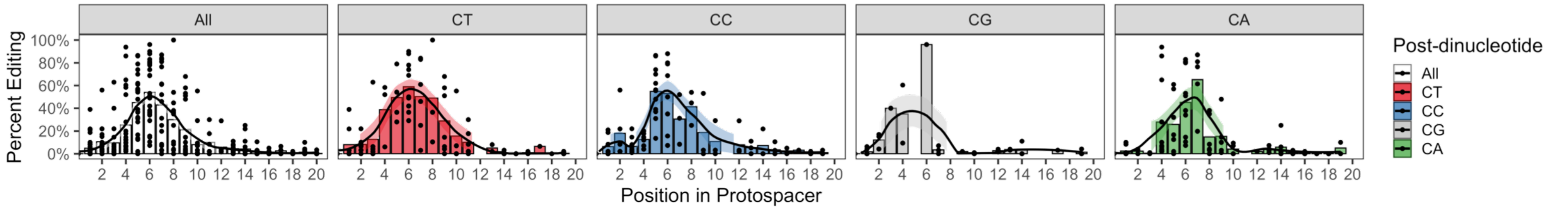


d

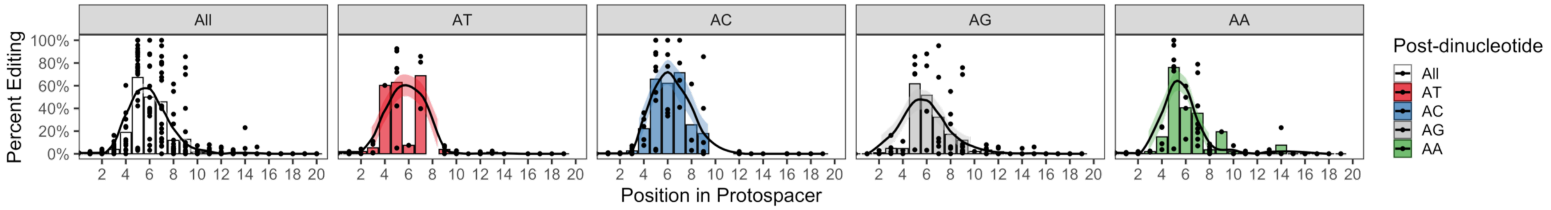


e

| PostDinucleotide | Average Editing | Std. Error | t value | Pr(>\|t\|) |
|---|---|---|---|---|
| All | 17.4% | 1.45% | 12.00 | 0.0000 |
| CT | 21.5% | 2.71% | 7.92 | 0.0000 |
| CC | 19.6% | 2.71% | 7.22 | 0.0000 |
| CA | 8.5% | 4.38% | 1.93 | 0.0534 |
| CG | 17.0% | 2.71% | 6.24 | 0.0000 |

f

| PostDinucleotide | Average Editing | Std. Error | t value | Pr(>\|t\|) |
|---|---|---|---|---|
| All | 12.6% | 1.34% | 9.35 | 0.00e+00 |
| AT | 10.7% | 3.24% | 3.29 | 1.04e-03 |
| AC | 20.1% | 2.84% | 7.10 | 0.00e+00 |
| AA | 9.0% | 2.35% | 3.82 | 1.46e-04 |
| AG | 13.5% | 2.73% | 4.94 | 9.00e-07 |

**Supplementary Figure 5.** Dinucleotide context dependencies of rAPOBEC1-BE4 and TadA$^{WT}$-TadA$^{Evo}$-ABE7.10. (a) Summary of linear model of rAPOBEC1-BE4 editing efficiency as a function of pre-dinucleotide context. (b) Summary of linear model of TadA$^{WT}$-TadA$^{Evo}$-ABE7.10 editing efficiency as a function of pre-dinucleotide context. (c) Distribution of APOBEC1-BE4 editing efficiency across the protospacer by post-dinucleotide context. (d) Distribution of TadA$^{WT}$-TadA$^{Evo}$-ABE7.10 editing efficiency across the protospacer by post-dinucleotide context. Note that distributions are not as smooth as pre-dinucleotide context (Fig. 4). (e) Summary of linear model of rAPOBEC1-BE4 editing efficiency as a function of post-dinucleotide context. (f) Summary of linear model of TadA$^{WT}$-TadA$^{Evo}$-ABE7.10 editing efficiency as a function of post-dinucleotide context.

### Figure S6

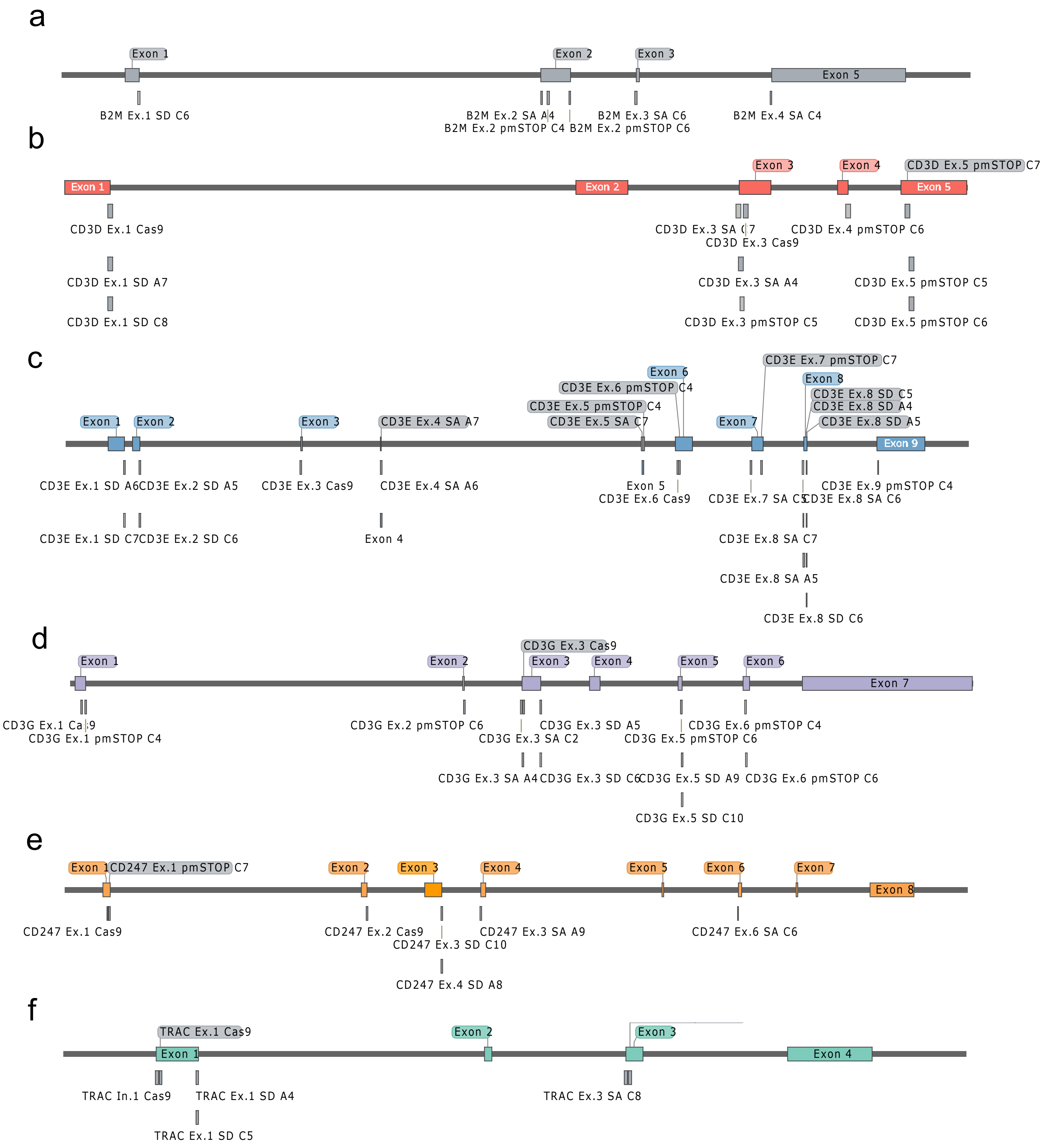


**Supplementary Figure 6.** Mapping of sgRNAs used in this work to the genomic loci of (a) B2M, (b) CD3D, (c) CD3E, (d) CD3G, (e) CD247, (f) TRAC. TRBC1 and TRBC2 were omitted from the BE-splice screen due to the inability to design single BE-splice sgRNAs to target both paralogs simultaneously.

Figure S7

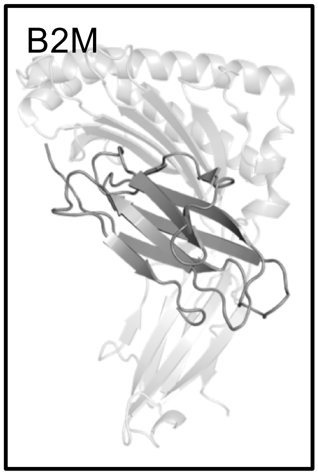


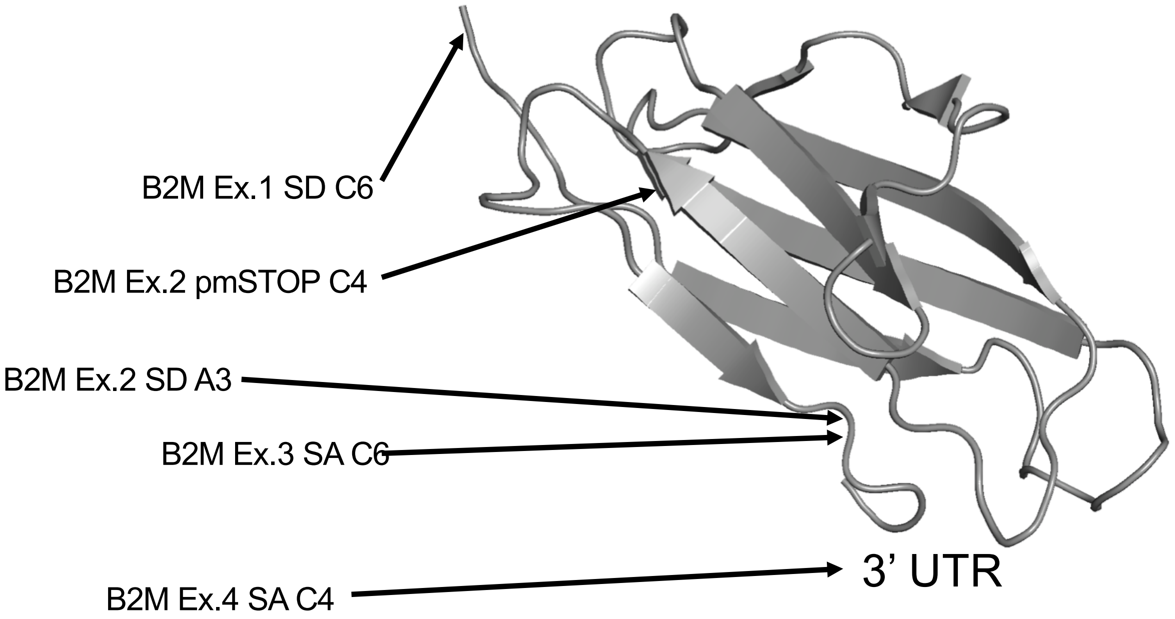


**Supplementary Figure 7**. Mapping of sgRNAs to B2M (β2M) protein structure (PDB 10GA).

### Figure S8

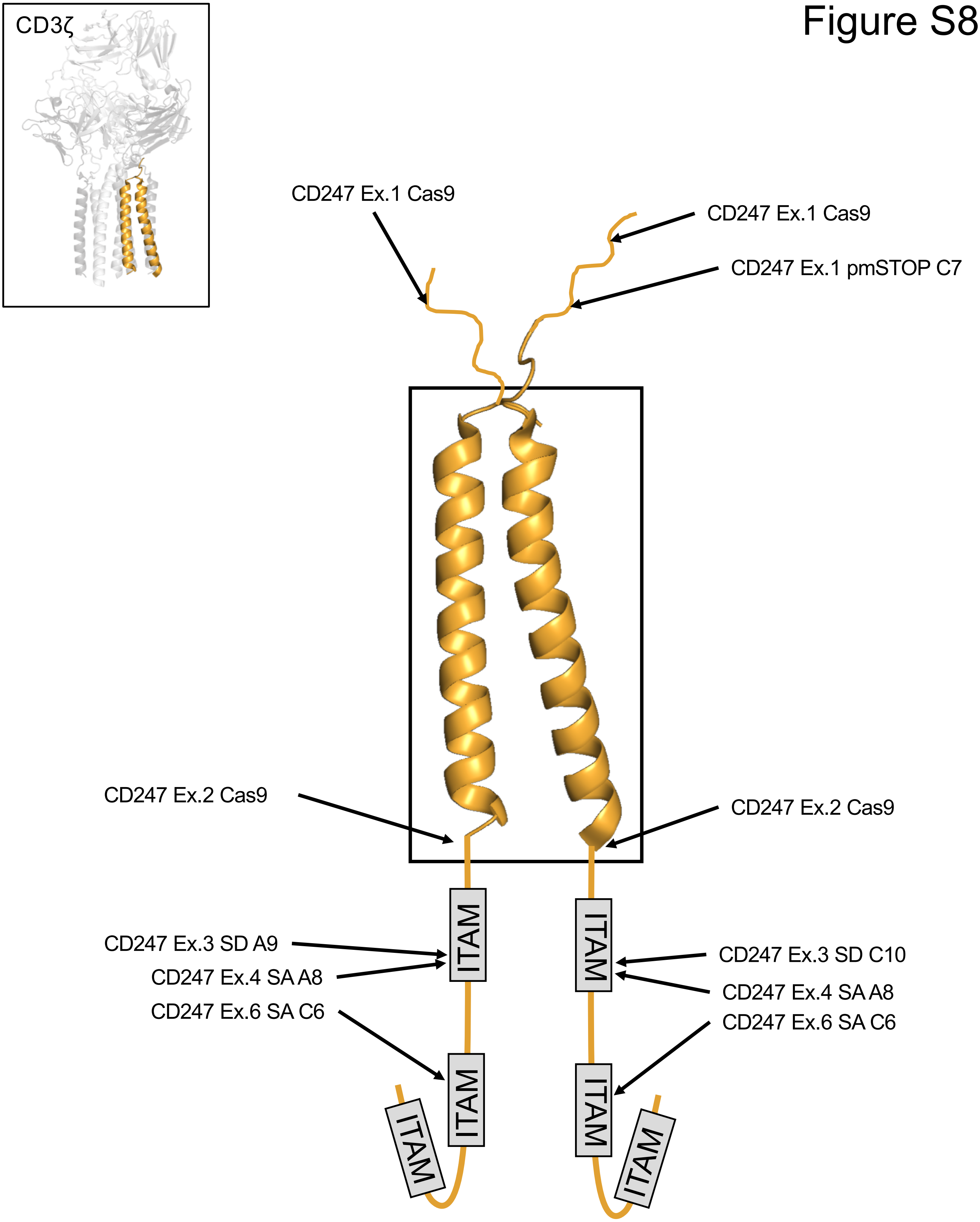


**Supplementary Figure 8.** Mapping of sgRNAs to CD247 (CD3ζ) protein structure (PDB 6JXR).

### Figure S9

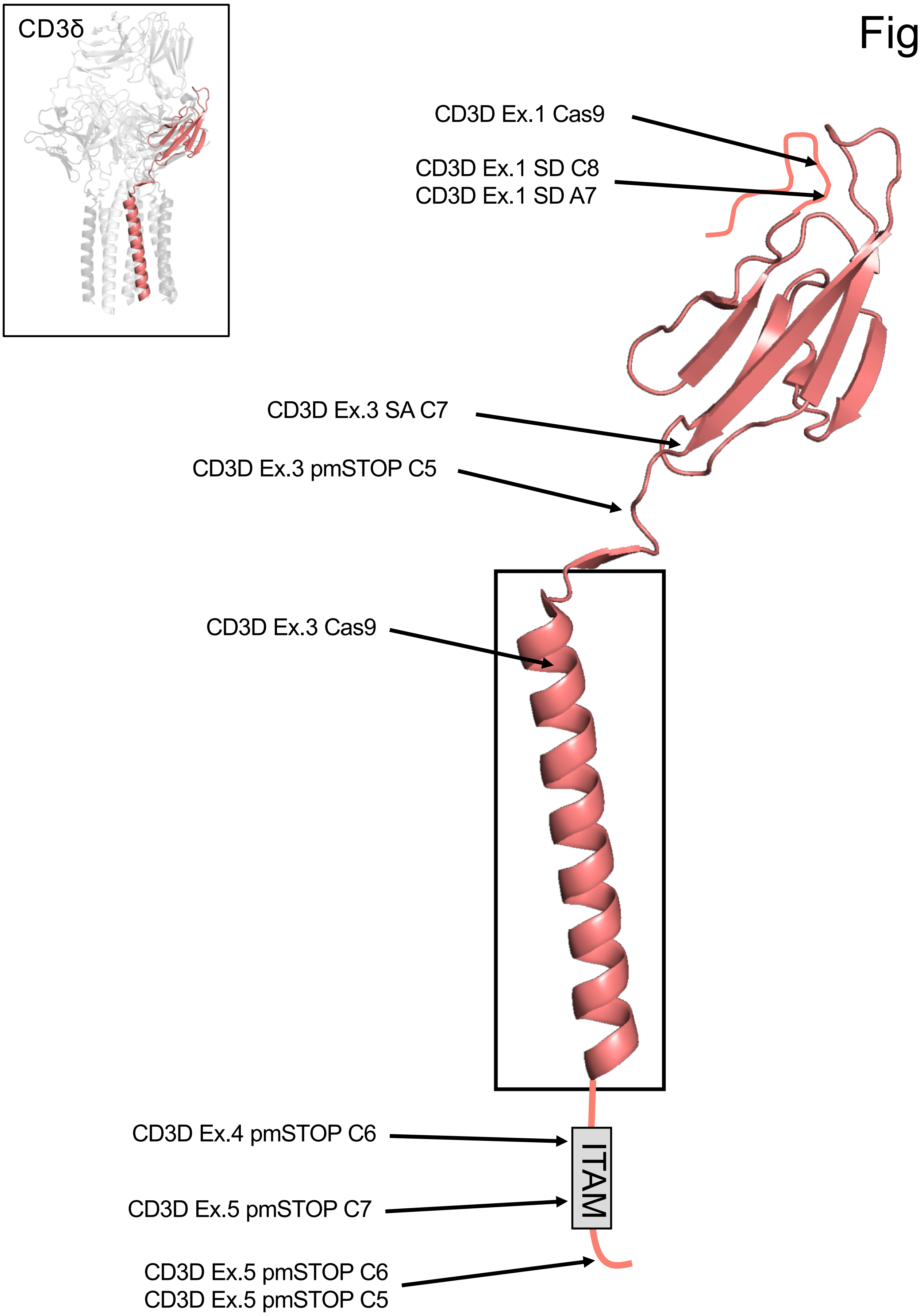


**Supplementary Figure 9.** Mapping of sgRNAs to CD3D protein structure (PDB 6JXR).

Figure S10

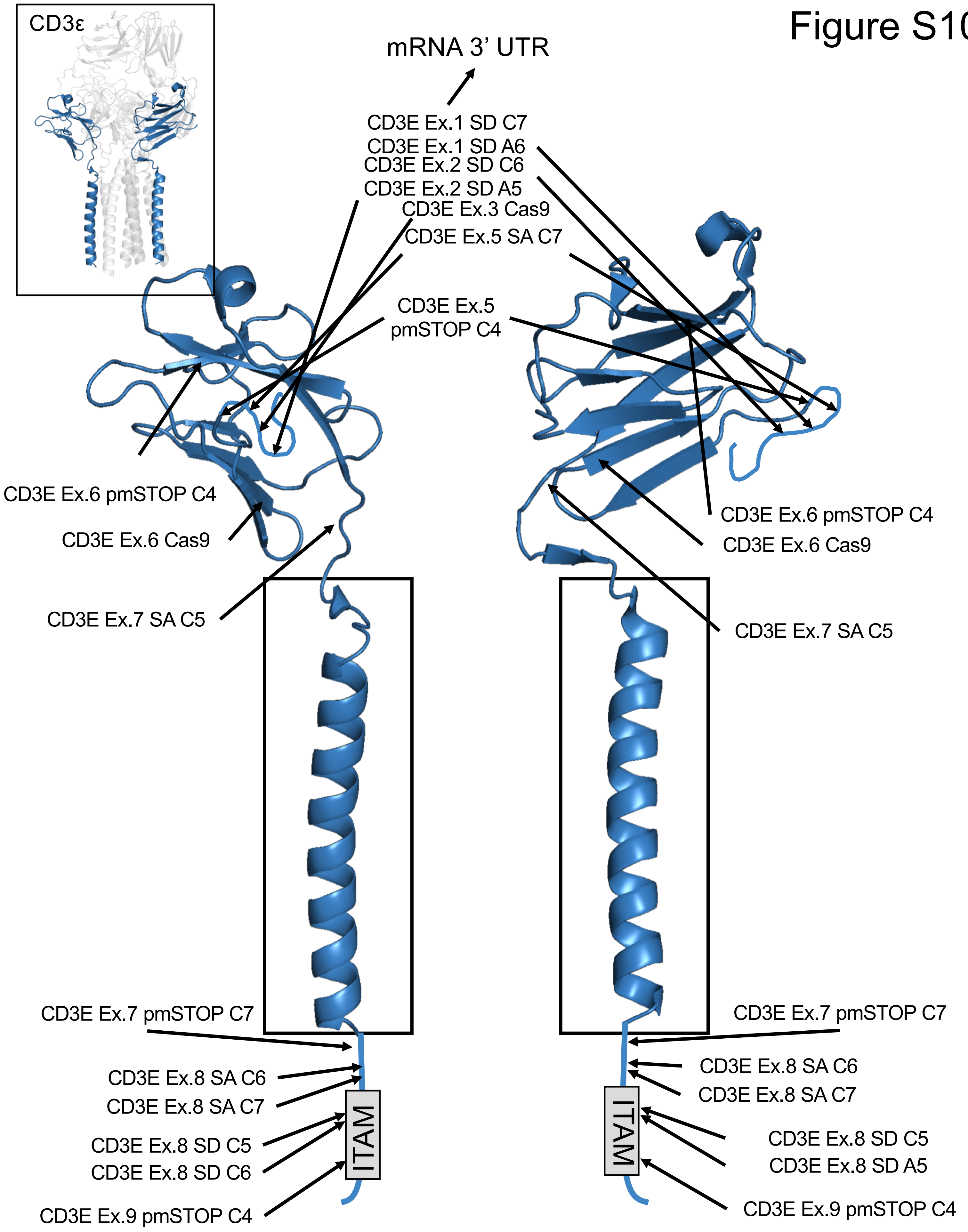


**Supplementary Figure 10.** Mapping of sgRNAs to CD3E protein structure (PDB 6JXR).

Figure S11

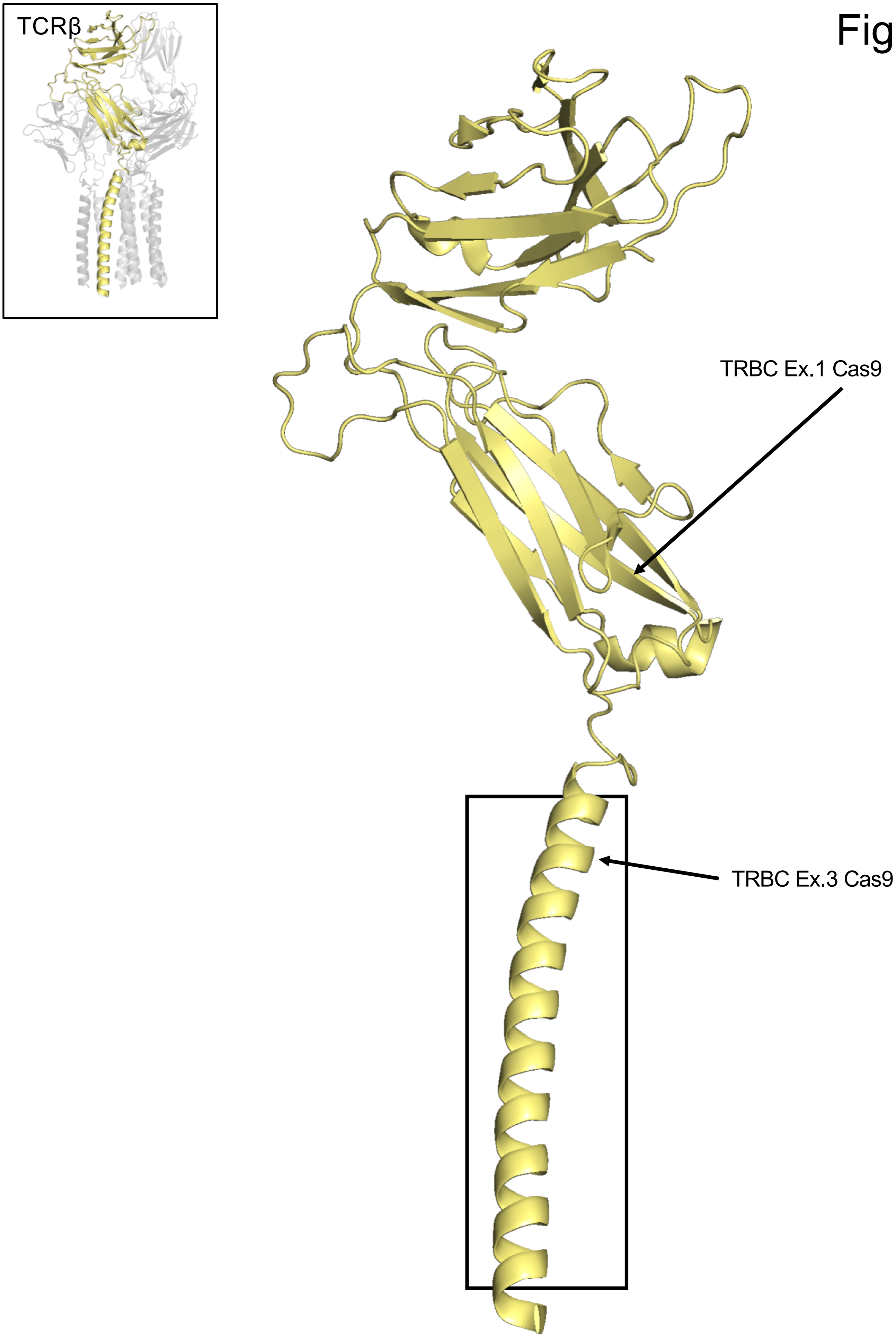


**Supplementary Figure 11.** Mapping of sgRNAs to TRBC (TCRβ) protein structure (PDB 6JXR).

### Figure S12

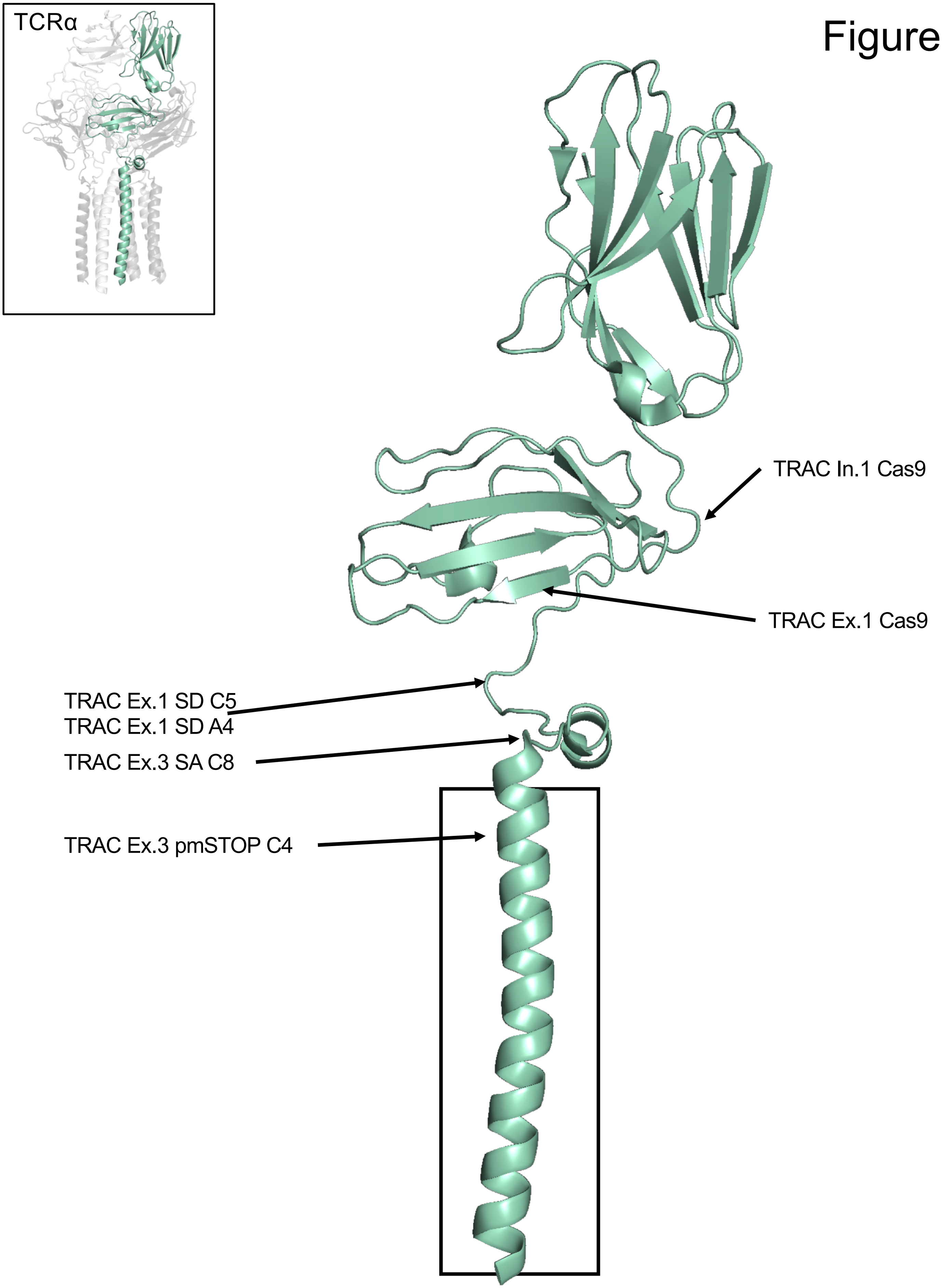


**Supplementary Figure 12.** Mapping of sgRNAs to TRAC (TCRα) protein structure (PDB 6JXR).

### Figure S13

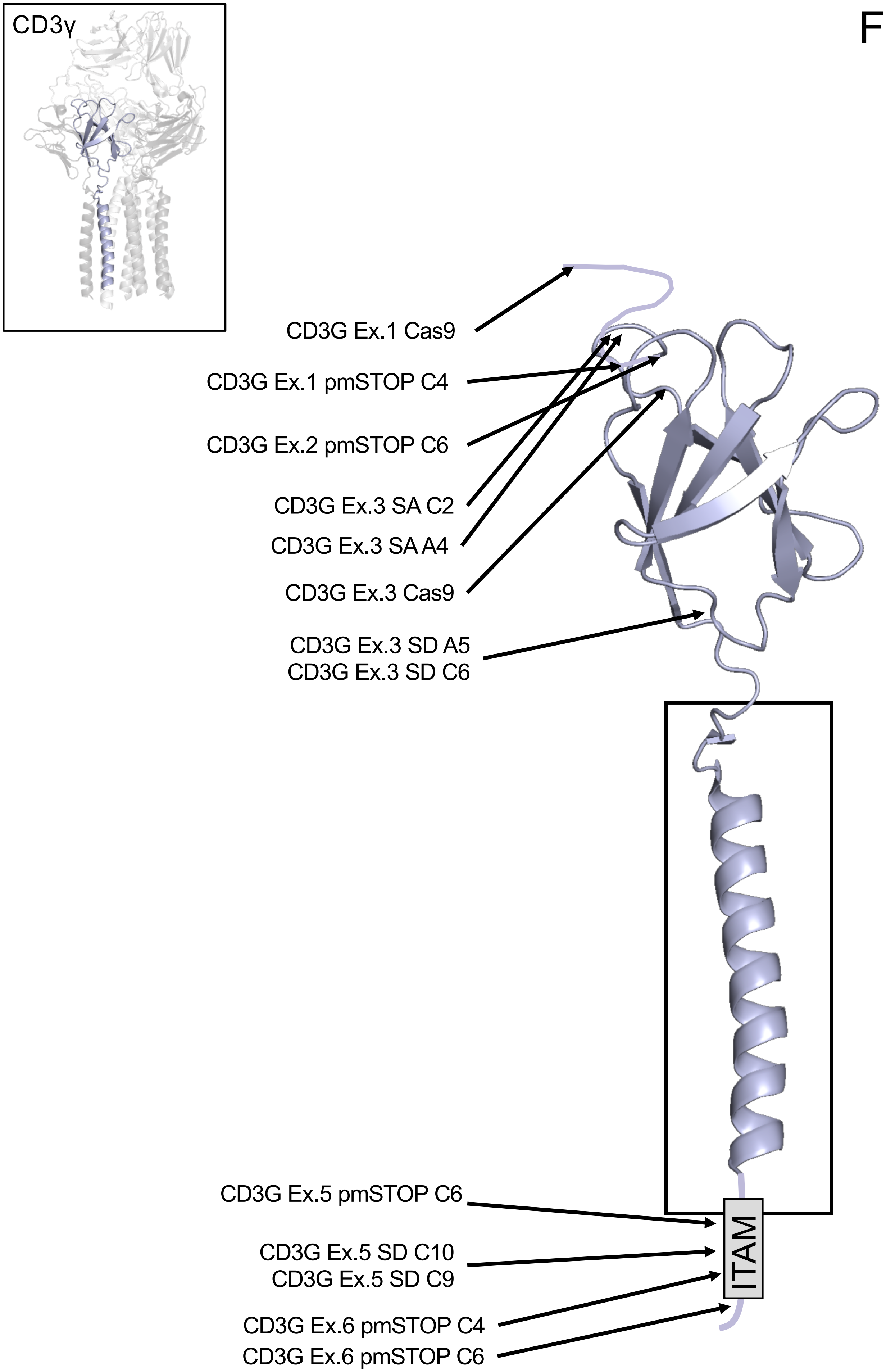


**Supplementary Figure 13.** Mapping of sgRNAs to CD3G protein structure (PDB 6JXR).
