## Additional file 1 for "CRISPR-Cas9 cytidine and adenosine base editing of splice-sites mediates highly-efficient disruption of proteins in primary cells"

```
### Figures for SpliceR

###############################################################################################
# Copyright (C) 2018-2019 Mitchell Kluesner
#
# This file is part of SpliceR Project
#
# Please only copy and/or distribute this script with proper citation of
# Splicer publication
###############################################################################################

##### LIBRARIES #####
library(tidyverse)
library(magrittr)
library(scales)
library(gridExtra)
library(googlesheets)
library(ggseqlogo)

####### Main Figures #######

##### FIGURE 1 #####
### Figure 1A
# Diagram of base editor

### Figure 1B
# Diagram of BE-splice method

# Table of practicality of approach

# Color palettes
colors5 = c("#d7191c", "#fdae61", "khaki", "#abd9e9", "#2c7bb6")
colors = c("NGA" = "#66c2a5", "NGT" = "#fc8d62", "NGC" = "#8da0cb", "NGG" = "#e78ac3", "NGN" =
"#b3b3b3")

## Define functions
len_uni = function(x){length(unique(x))}

as.percent = function(x, decimal_place = 4){paste0(round(x, decimal_place)*100, "%")}

guides =
read_tsv("/Users/kluesner/Desktop/Research/spliceR/submissions/additional_files/protein_guides_1-
95563_2018.09.06.tsv") %>%
  # Convert the position_score to position in the molecule
  mutate(position_score = 1-position_score) %>%
  # Remove any empty entries that  did not produce any guides
  filter(!is.na(be)) %>%
  # Remove the version number from the Ensembl transcript ID to be paired with other dataset in later
analysis
  tidyr::separate(., col = id, into = c("tmp_id"), sep = '[.]', remove = FALSE) %>%
  # convert BE to include if it is targetable by BE3 or ABE
  mutate(be = {ifelse(splice_site == "donor", "BE3 or ABE", be)}) %>%
  # Make a generic PAM class for each pam
  mutate(pam_class = sub("^.", "N", pam))

### Establish genes used for guide screen
genes =
read_tsv("/Users/kluesner/Desktop/Research/spliceR/submissions/additional_files/protein_genes_1-
97713_2018.09.06.tsv") %>%
  # Remove the version number from the Ensembl transcript ID to be paired with other dataset in later
analysis
  tidyr::separate(., col = id, into = c("tmp_id"), sep = '[.]', remove = FALSE)

### Establish various IDs for different parts of the analysis
# Ensembl transcript IDs
ids =
read_tsv("/Users/kluesner/Desktop/Research/spliceR/submissions/additional_files/protein.txt")$ENST

# Ensembl gene IDs
gene_ids = genes$gene_id %>% unique()
```

```
# Ensembl transcript IDs a that are non-single exon, single isoform
nsesi_ids = genes %>%
  group_by(id) %>%
  summarize(number = length(id)) %>%
  filter(number > 1) %>%
  .$id

# Ensembl gene IDs a that are non-single exon, single isoform
nsesi_gene_ids = genes %>%
  group_by(gene_id) %>%
  summarize(number = length(gene_id)) %>%
  filter(number > 1) %>%
  .$gene_id

## Basic stats
    # Percent of all transcript ids that are non-single-exon-single-isoform (NSESI)
    perc_nsesi_ids = length(nsesi_ids)/length(ids)

    # Percent of all gene ids that are NSESI
    perc_nsesi_gene_ids = length(nsesi_gene_ids)/length(gene_ids)

    ## Transcripts
    # Number of transcript ids that yielded a gRNA that allows it to be targetable
    n_targettable_ids = (ids %in% guides$id) %>% sum()

    # Percent of total transcript ids that are targetable
    perc_targettable_ids = n_targettable_ids/length(ids)

    # Number of NSESI transcript ids of all of the ensembl transcript ids
    n_nsesi_targettable_ids = (nsesi_ids %in% guides$id) %>% sum()

    # Percent of NSESI transcript ids that are targetable (i.e. have potential guides)
    perc_nsesi_targettable_ids = n_targettable_ids/length(nsesi_ids)

    ## Genes
    # Number of genes that were found to have guides to them, i.e. targetable
    n_targettable_genes = gene_ids %in% guides$gene_id %>% sum()

    # percent of all genes that were found to have a gRNA to them
    perc_targettable_genes = n_targettable_genes/length(gene_ids)

    # Number of ensembl gene ids that are NSESI
    n_nsesi_targettable_genes = nsesi_gene_ids %in% guides$gene_id %>% sum()

    # Percent of NSESI gene ids that were found to have a gRNA to them
    perc_nsesi_targettable_genes = n_nsesi_targettable_genes/length(nsesi_gene_ids)



# Scoring was defined based on Komor et al., 2016 and Gaudelli et al., 2017
figure_1C = tibble(`Protein Coding` = c("Transcripts", "Genes"),
                   Number = c(n_targettable_ids, n_targettable_genes),
                   `Percent of NSESI` = c(perc_nsesi_targettable_ids, perc_nsesi_targettable_genes)
%>% signif(4) %>% scales::percent(),
                   `Percent of Total` = c(perc_targettable_ids, perc_targettable_genes) %>% signif(4)
%>% scales::percent()
)

### Figure 1D
# Cumulative percent of targetable genes as a function of the earliest targetable splice-site

figure_1D = figure_1_data %>%
  ggplot(aes(x = number_of_guides, color = PAM)) +
  stat_ecdf(geom = "step", pad = FALSE, lwd = 1.5, alpha = 0.9) +
  scale_y_continuous(labels = scales::percent_format(), limits = c(0, 1)) +
  scale_x_log10() +
  annotation_logticks(sides = "b") +
  ylab("Cumulative % of targetable genes") +
  xlab("Number of BE-Splice gRNAs per gene") +
```

```
  coord_cartesian(xlim = c(0.9999, max(figure_1_data$number_of_guides)), ylim = c(-0.005, 1.005),
expand = F) +
  theme_bw(base_size = 24) +
  theme(panel.border = element_rect(colour = "black", fill=NA),
        aspect.ratio = 1,
        legend.position = c(0.825,  0.6),
        legend.background = element_rect(fill = "white", color = "black"),
        plot.margin = unit(c(1,1,1,1), "cm")) +
  labs(color = "PAM") +
  scale_color_manual(values = colors) +
  annotate(geom = "segment",  x = number50, xend = number50, y = 0, yend = 0.5, lwd = 1, linetype =
"dashed") +
  annotate(geom = "segment",  x = 0, xend = number50, y = 0.5, yend = 0.5, lwd = 1, linetype =
"dashed") +
  annotate(geom = "text", size = 8, x = 10^(0.14*log10(max(figure_1_data$number_of_guides))), y =
0.55, label = paste0(number50, " guides"))

### Figure 1D
# Cumulative percent of targetable genes a sa function of the number of BE-splice gRNAs per gene
figure_1_data = guides %>%
  mutate(NGN = "NGN")  %>%
  gather(type, PAM, c("pam_class", "NGN")) %>%
  group_by(gene_id, PAM) %>%
  dplyr::summarise(earliest_guide = min(position_score), number_of_guides = length(position_score))
%>%
  mutate(PAM = factor(PAM, levels = c("NGG", "NGA", "NGT", "NGC", "NGN")))

guide50 = figure_1_data %>% filter(PAM == "NGN") %>% .$earliest_guide %>% median
number50 = figure_1_data %>% filter(PAM == "NGN") %>% .$number_of_guides %>% median

## make figure 1E
figure_1E = figure_1_data %>%
  ggplot(aes(x = earliest_guide, color = PAM)) +
  stat_ecdf(geom = "step", pad = FALSE, lwd = 1.5, alpha = 0.8) +
  scale_y_continuous(labels = scales::percent_format(), limits = c(0, 1)) +
  scale_x_continuous(limits = c(0,1.2), breaks = seq(0, 1.2, 0.2)) +
  ylab("Cumulative % of targetable genes") +
  xlab("Earliest BE-Splice targetable splice-site\n(relative position in mRNA)") +
  coord_cartesian(xlim = c(0, 1.0), ylim = c(0, 1), expand = F) +
  theme_bw(base_size = 24) +
  theme(panel.border = element_rect(colour = "black", fill=NA),
        aspect.ratio = 1,
        legend.position = c(0.825,  0.6),
        legend.background = element_rect(fill = "white", color = "black"),
        plot.margin = unit(c(1,1,1,1), "cm")) +
  labs(color = "PAM") +
  scale_color_manual(values = colors) +
  annotate(geom = "segment",  x = guide50, xend = guide50, y = 0, yend = 0.5, lwd = 1, linetype =
"dashed") +
  annotate(geom = "segment",  x = 0, xend = guide50, y = 0.5, yend = 0.5, lwd = 1, linetype =
"dashed") +
  annotate(geom = "text", size = 8, x = 0.3, y = 0.44, label = paste0(as.percent(guide50), "
into\nmRNA"))

##### FIGURE 2 #####
### Figure 2A
# Diagram of complex

### Figure 2B
# Diagram of screen rationale

### Figure 2C
# Genetic and protein KO from STE2

gene_colors = c(B2M = "#d9d9d9", TRAC = '#8dd3c7', TRBC = '#ffffb3', CD3G = '#bebada', CD3D =
'#fb8072',CD3E = '#80b1d3',CD247 = '#fdb462', AAVS1  = "#b3de69")

experimental_data =
read_tsv("/Users/kluesner/Desktop/Research/spliceR/submissions/additional_files/compiled_STE_data.tsv")
 %>%
```

```
  mutate(Indel = Indel/100) %>%
  mutate(Edit = {ifelse(Enzyme == "BE4", `T`, `G`)/100}) %>%
  mutate(Protein_Loss = 1-Flow) %>%
  mutate(Gene = factor(Gene, levels =  c("TRAC", "TRBC", "CD3D", "CD3E", "CD3G", "CD247", "B2M",
"AAVS1"))) %>%
  mutate(Edit = {ifelse(Enzyme == "Cas9", Indel, Edit)}) %>%
  mutate(Enzyme_Motif = paste0(Enzyme, " ", Motif)) %>%
  mutate(Enzyme_Motif = factor(Enzyme_Motif, levels = c("BE4 SD", "BE4 SA", "BE4 pmSTOP", "ABE7.10
SD", "ABE7.10 SA", "Cas9 Control"))) %>%
  mutate(Exon_Class = {ifelse(Exon == 1, "First",
                              ifelse(Exon == 2, "Second",
                                     ifelse(Exon == N_Exons, "Last",
                                            ifelse(Exon == N_Exons - 1, "Second-to-last",
"Middle"))))}) %>%
  mutate(Exon_Class = factor(Exon_Class, levels = c("First", "Second", "Middle", "Second-to-last",
"Last"))) %>%
  mutate(Exon_Group = {ifelse(Exon < N_Exons - 1, Exon,
                              ifelse(Exon == N_Exons - 1, "Second-to-last",
                                     ifelse(Exon == N_Exons, "Last", "NA")))}) %>%
  mutate(Exon_Group = factor(Exon_Group, levels = c(1:7, "Second-to-last", "Last"))) %>%
  mutate(Enzyme = factor(Enzyme, levels = c("BE4", "ABE7.10", "Cas9"))) %>%
  mutate(Motif = factor(Motif, levels = c("SA", "SD", "pmSTOP")))

figure_2C = experimental_data %>%
  filter(Experiment == "STE2") %>%
  filter(Gene != "AAVS1") %>%
  dplyr::rename(`Protein Loss` = Protein_Loss) %>%
  gather(metric, value, c("Protein Loss", "Indel")) %>%
  ggplot(aes(x = Guide_Name, y = value, fill = Gene)) +
  geom_bar(stat = "summary", fun.y = "mean", color  = 'black') +
  geom_point() +
  scale_y_continuous(limits = c(0,1), breaks = seq(0,1,0.2), labels = scales::percent_format()) +
  scale_fill_manual(values = gene_colors) +
  facet_grid(cols = vars(Gene), rows = vars(metric), space = "free_x", scale = "free_x") +
  ylab("Percent knockout") +
  xlab("") +
  theme_bw(base_size = 18) +
  theme(axis.text.x = element_text(hjust = 1, angle = 45),
        panel.grid.major = element_blank(), panel.grid.minor = element_blank())

### Figure 2 statistics
figure_2_statistics = experimental_data %>%
  filter(Experiment == "STE2") %>%
  group_by(Experiment) %>%
  dplyr::summarise(mean_indel = mean(Edit), mean_protein_loss = mean(Protein_Loss),
                   sd_indel = sd(Edit), sd_protein_loss = sd(Protein_Loss)) %>%
  ungroup() %>%
  dplyr::select(-Experiment) %>%
  mutate_all(~signif(.,digits= 3) %>% scales::percent()) %>%
  mutate(Experiment = "STE2") %>%
  dplyr::select(Experiment, everything())


##### FIGURE 3 #####
### Figure 3A
# Genetic and protein KO bargraphs
figure_3A = experimental_data %>%
  filter(Experiment != "STE2") %>%
  mutate(Enzyme_factor = {ifelse(Enzyme ==  "BE4", 1, 2)}) %>%
  dplyr::rename(`Protein Loss` = Protein_Loss) %>%
  gather(metric, value, c("Protein Loss", "Edit")) %>%
  ggplot(aes(x = reorder(Guide_Name, -value)  %>% reorder(., Enzyme_factor), y = value, fill = Gene,
shape  = Enzyme)) +
  geom_bar(stat = "summary", fun.y = "mean", color  = 'black') +
  geom_point(size = 3, aes(color = Enzyme)) +
  scale_color_manual(values = c("BE4" = "black", "ABE7.10" = "darkred")) +
  scale_y_continuous(limits = c(0,1), breaks = seq(0,1,0.2), labels = scales::percent_format()) +
  scale_fill_manual(values = gene_colors) +
  facet_grid(cols = vars(Gene), rows = vars(metric), space = "free", scale = "free") +
  ylab("") +
```

```
  xlab("") +
  theme_bw(base_size = 24) +
  theme(axis.text.x = element_text(hjust = 1, angle = 60),
        panel.grid.major = element_blank(), panel.grid.minor = element_blank())

### Figure 3A statistics
figure_3A_statistics = experimental_data %>%
  filter(Experiment != "STE2") %>%
  mutate(Experiment = "STE3 and STE4") %>%
  group_by(Experiment) %>%
  dplyr::summarise(mean_edit = mean(Edit), mean_protein_loss = mean(Protein_Loss),
                   sd_edit = sd(Edit), sd_protein_loss = sd(Protein_Loss),
                   min_edit = min(Edit), max_edit = max(Edit),
                   # median_edit = median(Edit), median_protein_loss = median(Protein_Loss),
                   min_protein_loss = min(Protein_Loss), max_protein_loss = max(Protein_Loss)) %>%
  ungroup() %>%
  dplyr::select(-Experiment) %>%
  mutate_all(~signif(.,digits= 3) %>% scales::percent()) %>%
  mutate(Experiment = "STE3 and STE4") %>%
  dplyr::select(Experiment, everything())

### Figure 3B
# Genetic boxplot

set.seed(1)
figure_3B = experimental_data %>%
  filter(Experiment != "STE2") %>%
  # filter(Domain != "Intracellular") %>% # Conditional look at what happens when you remove all
intracellular premature stop codons
  ggplot(aes(x = Motif, y = Edit, fill = Enzyme)) +
  geom_boxplot(outlier.alpha  = 0) +
  geom_point(pch = 21,position = position_jitter(0.1), alpha= 0.7, fill = "darkgrey", size = 3) +
  xlab("") +
  ylab("Editing of target base") +
  #scale_color_manual(values = c("acceptor"= "#a6cee3", "donor" = "#1f78b4", "pmSTOP" = "#b2df8a")) +
  scale_fill_manual(values = c("BE4" = "#1f78b4", "ABE7.10" = "#b2df8a")) +
  scale_y_continuous(limits = c(-0.01, 1.2), labels = scales::percent_format(), breaks = seq(0,1,0.2))
+
  # scale_x_discrete(labels = c("SA", "SD", "pmSTOP")) +
  #geom_bar(stat = "summary", fun.y = "median")  +
  facet_grid(cols = vars(Enzyme), scales = 'free_x', space = "free") +
  theme_bw(base_size = 24) +
  theme(panel.grid.major = element_blank(), panel.grid.minor = element_blank())

### Figure 3C
# Protein boxplot

set.seed(1)
figure_3C = experimental_data %>%
  filter(Experiment != "STE2") %>%
  # filter(Domain != "Intracellular") %>% # Conditional look at what happens when you remove all
intracellular premature stop codons
  ggplot(aes(x = Motif, y = Protein_Loss, fill = Enzyme)) +
  geom_boxplot(outlier.alpha  = 0) +
  # geom_bar(stat = "summary", fun.y = "mean") +
  geom_point(pch = 21,position = position_jitter(0.1), alpha= 0.7, fill = "darkgrey", size = 3) +
  xlab("") +
  ylab("Loss in surface expression") +
  #scale_color_manual(values = c("acceptor"= "#a6cee3", "donor" = "#1f78b4", "pmSTOP" = "#b2df8a")) +
  scale_fill_manual(values = c("BE4" = "#1f78b4", "ABE7.10" = "#b2df8a")) +
  scale_y_continuous(limits = c(-0.01, 1.2), labels = scales::percent_format(), breaks = seq(0,1,0.2))
+
  # scale_x_discrete(labels = c("SA", "SD", "pmSTOP")) +
  #geom_bar(stat = "summary", fun.y = "median")  +
  facet_grid(cols = vars(Enzyme), scales = 'free_x', space = "free") +
  theme_bw(base_size = 24) +
  theme(panel.grid.major = element_blank(), panel.grid.minor = element_blank())

### Figure 3D
# Correlation between editing and KO across STE2, STE3 and STE4
```

```
# Define function for correlation
correlation = function(x, y){cor.test(x = x, y = y)$estimate}
correlation_pvalue = function(x, y){cor.test(x = x, y = y)$p.value}

motif_cor_data = experimental_data %>%
  group_by(Enzyme_Motif) %>%
  dplyr::summarize(cor = signif(correlation(Edit, Protein_Loss), 3),
                   pvalue = signif(correlation_pvalue(Edit, Protein_Loss), 3)) %>%
  ungroup() %>%
  # mutate(Enzyme_Motif = paste0(Enzyme, " ", Motif)) %>%
  #mutate(label = paste0())
  mutate(var = Enzyme_Motif) %>%
  mutate(x  = 0.25, y = 0.75) %>%
  mutate(cor = paste0("r = ", cor)) %>%
  mutate(pvalue = paste0("p-value = ", pvalue))

figure_3D = experimental_data %>%
  # mutate(Enzyme_Motif = factor()) $%>
  ggplot(aes(x  = Edit, y = Protein_Loss, fill = Enzyme_Motif)) +
  geom_abline(slope = 1, intercept = 0, linetype = "dashed") +
  geom_smooth(method = "lm", color = "black", aes(fill = NULL)) +
  geom_point(pch = 21) +
  scale_y_continuous(labels = scales::percent_format(), limits = c(0, 1)) +
  scale_x_continuous(labels = scales::percent_format(), limits = c(0, 1)) +
  theme_bw(base_size = 18) +
  geom_label(data = motif_cor_data, aes(x = x, y = y, label = cor, fill = NULL)) +
  xlab("Genetic Editing") +
  ylab("Protein Loss") +
  facet_wrap(.~Enzyme_Motif) +
  theme(aspect.ratio = 1,
        axis.text.x = element_text(hjust = 1, angle = 45),
        legend.position = "none",
        panel.grid.major = element_blank(),
        panel.grid.minor = element_blank()
        )


### FIGURE 3 STATISTICS ###
### CBE vs. ABE
# equal variance
experimental_data %>% filter(Experiment != "STE2") %>% filter(Motif != "pmSTOP") %>% var.test(Edit ~
Enzyme, data =  .)
experimental_data %>% filter(Experiment != "STE2") %>% filter(Motif != "pmSTOP") %>% t.test(Edit ~
Enzyme, data =  ., var.equal = T)

experimental_data %>% filter(Experiment != "STE2") %>% filter(Motif != "pmSTOP") %>%
var.test(Protein_Loss ~ Enzyme, data =  .)
experimental_data %>% filter(Experiment != "STE2") %>% filter(Motif != "pmSTOP") %>%
t.test(Protein_Loss ~ Enzyme, data =  ., var.equal = T)

### SD vs.SA
## Across ABE and CBE
# Equal variance
# No significant difference in editing between SDs and SAs across ABEs and CBEs
experimental_data %<>% mutate(Motif = as.character(levels(Motif)[Motif]))

experimental_data %>% filter(Experiment != "STE2") %>% filter(Motif != "pmSTOP") %>% var.test(Edit ~
Motif, data =  .)
# One the cusp of being significant
experimental_data %>% filter(Experiment != "STE2") %>% filter(Motif != "pmSTOP") %>% t.test(Edit ~
Motif, data =  ., var.equal = T)


### CBE
### Statistics
CBE_editing_mean = experimental_data %>% filter(Enzyme == "BE4") %>% .$Edit %>% mean(., na.rm = T)
CBE_editing_sd = experimental_data %>% filter(Enzyme == "BE4") %>% .$Edit %>% sd(., na.rm = T)

CBE_protein_mean = experimental_data %>% filter(Enzyme == "BE4") %>% .$Protein_Loss %>% mean(., na.rm
```

```
= T)
CBE_protein_sd = experimental_data %>% filter(Enzyme == "BE4") %>% .$Protein_Loss %>% sd(., na.rm = T)

# Equal variance of data
# Average editing between SD and SA among CBE is n.s.
experimental_data %>% filter(Motif != "pmSTOP" & Enzyme == "BE4") %>% var.test(Edit ~ Motif, data =
.)
experimental_data %>% filter(Motif != "pmSTOP" & Enzyme == "BE4") %>% t.test(Edit ~ Motif, data =  .,
var.equal = T)

# Equal variance of data

# Average protein loss between SD and SA among CBE is n.s.
experimental_data %>% filter(Motif != "pmSTOP" & Enzyme == "BE4") %>% var.test(Protein_Loss ~ Motif,
data =  .)
experimental_data %>% filter(Motif != "pmSTOP" & Enzyme == "BE4") %>% t.test(Protein_Loss ~ Motif,
data =  ., var.equal = T)

### ABE
### Statistics
ABE_editing_mean = experimental_data %>% filter(Enzyme == "ABE7.10") %>% .$Edit %>% mean(., na.rm = T)
ABE_editing_sd = experimental_data %>% filter(Enzyme == "ABE7.10") %>% .$Edit %>% sd(., na.rm = T)

ABE_protein_mean = experimental_data %>% filter(Enzyme == "ABE7.10") %>% .$Protein_Loss %>% mean(.,
na.rm = T)
ABE_protein_sd = experimental_data %>% filter(Enzyme == "ABE7.10") %>% .$Protein_Loss %>% sd(., na.rm
= T)

experimental_data %>%
  filter(Enzyme == "BE4" & Motif == "pmSTOP") %$%
  cor.test(x = Edit, y = Protein_Loss)

# Data does not have equal variance
# Average editing between SD and SA among ABE is significantly different
experimental_data %>% filter(Motif != "pmSTOP" & Enzyme == "ABE7.10") %>% var.test(Edit ~ Motif, data
=  .)
experimental_data %>% filter(Motif != "pmSTOP" & Enzyme == "ABE7.10") %>% t.test(Edit ~ Motif, data =
., var.equal = F)

# Data does not have equal variance
# Average protein loss between SD and SA among ABE is significantly different
experimental_data %>% filter(Motif != "pmSTOP" & Enzyme == "ABE7.10") %>% var.test(Protein_Loss ~
Motif, data =  .)
experimental_data %>% filter(Motif != "pmSTOP" & Enzyme == "ABE7.10") %>% t.test(Protein_Loss ~ Motif,
data =  ., var.equal = F)

### SD vs. SA vs. pmSTOP
# Just focus on CBE for fair comparison

# Equal variance of data
# Average editing between SD and pmSTOP is n.s.
experimental_data %>% filter(Motif != "SA" & Enzyme == "BE4") %>% var.test(Edit ~ Motif, data =  .)
experimental_data %>% filter(Motif != "SA" & Enzyme == "BE4") %>% t.test(Edit ~ Motif, data =  .,
var.equal = T)

# Variance of data is equal
# Average protein loss between SD and pmSTOP among CBE is significant
experimental_data %>% filter(Motif != "SA" & Enzyme == "BE4") %>% var.test(Protein_Loss ~ Motif, data
=  .)
experimental_data %>% filter(Motif != "SA" & Enzyme == "BE4") %>% t.test(Protein_Loss ~ Motif, data =
., var.equal = T)

# Just focus on CBE for fair comparison
# Equal variance of data
# Average editing between SA and pmSTOP is n.s.
# Although close
experimental_data %>% filter(Motif != "SD" & Enzyme == "BE4") %>% var.test(Edit ~ Motif, data =  .)
experimental_data %>% filter(Motif != "SD" & Enzyme == "BE4") %>% t.test(Edit ~ Motif, data =  .,
var.equal = T)
```

```
# Variance of data is equal, although close
# Average protein loss between SA and pmSTOP among CBE is n.s. significant but close
experimental_data %>% filter(Motif != "SD" & Enzyme == "BE4") %>% var.test(Protein_Loss ~ Motif, data
=  .)
experimental_data %>% filter(Motif != "SD" & Enzyme == "BE4") %>% t.test(Protein_Loss ~ Motif, data =
., var.equal = T)

### END STATISTICS ###


##### FIGURE 4 #####
### Define functions

flushOutData = function(df, base){
  # df = abe_data
  # base = "A"
  # Take a rowise approach
  # create df_a that has the cols Protospacer, Cell type, Paper, normalization, G_ave = 0, and G_norm
  df_a = df %>% dplyr::select(Protospacer)

  # create df_b data frame that has the cols Position, and Protospacer for each guide
  guides = df_a$Protospacer %>% unique()
  list_a = str_locate_all(guides, base)

  names(list_a) = guides
  vec_b = unlist(list_a)

  df_b = data.frame(Protospacer = names(vec_b), Position = vec_b) %>%
    mutate(Protospacer = gsub("[0-9]", "", Protospacer)) %>%
    distinct()

  # df_c = inner_join(df_a, df_b)
  df_c = inner_join(df_a, df_b) %>% distinct()

  # df_d = remove any rows from df_c that are in the data, i.e remove the filler rows for already
documented values
  df_d = anti_join(df_c, df) %>%
    mutate(Edit_ave = 0, Edit_norm = 0)

  # bind_cols(data, df_d)
  df_e = bind_rows(df_d, df)

  # recalculate the motifs
  final_data = df_e %>%
    mutate(Trinucleotide = substr(Protospacer, Position-1, Position+1)) %>%
    mutate(Dinucleotide = substr(Protospacer, Position-1, Position)) %>%
    mutate(Dinucleotide = factor(Dinucleotide, levels = paste0(c("T", "C", "A", "G"), base))) %>%
    mutate(PostDinucleotide = substr(Protospacer, Position, Position+1)) %>%
    mutate(PostDinucleotide = factor(PostDinucleotide, levels = paste0(base, c("T", "C", "G", "A"))))
%>%
    mutate(TrinucleotideClass = gsub("[A|G]", "R", Trinucleotide) %>% gsub("[T|C]", "Y", .))

  filler_data = data.frame(Dinucleotide = c(rep(paste0(c("A", "T", "C", "G"), base), 4),
rep(paste0(c("A", "T", "C", "G"), base), 4)),
                           PostDinucleotide = c(rep(paste0(base, c("A", "T", "C", "G")), 4),
rep(paste0(base, c("A", "T", "C", "G")), 4)),
                           Edit_norm = rep(rep(0, 4), 4), Position = c(rep(rep(1, 4), 4), rep(rep(0,
4), 4)))

  output_data = bind_rows(final_data, filler_data)

  final_output_data = bind_rows(
    output_data %>% mutate(Dinucleotide = "All", PostDinucleotide = "All"),
    output_data
    ) %>%
    mutate(Dinucleotide = factor(Dinucleotide, levels = c("All", paste0(c("T", "C", "A", "G"),
base)))) %>%
    mutate(PostDinucleotide = factor(PostDinucleotide, levels = c("All", paste0(base, c("T", "C", "G",
"A")))))
```

```
  return(final_output_data)
}

### Figure 4A
### CBE base editing by position

## Load data
cbe_data =
read_tsv("/Users/kluesner/Desktop/Research/spliceR/submissions/additional_files/cbe_meta_data.tsv")
%>%

  mutate(Protospacer = toupper(Protospacer)) %>%
  mutate(Edit = `T`) %>%

  # Establish the maximum edit for each cell type in each paper
  group_by(`Cell type`, Paper) %>%
  mutate(paper_max = max(Edit)) %>%
  ungroup %>%

  # Normalize the edits to the max edit observed in the paper
  mutate(Edit_norm = Edit/paper_max) %>%

  # For each position in each unique guide, calculate the average normalized edit
  group_by(Position, Protospacer) %>%
  dplyr::summarise(Edit_norm = mean(Edit_norm)) %>%
  ungroup() %>%
  arrange(Protospacer)


### Figure 4A
# CBE base editing by position, by dinucleotide
# Need to break up in the actual figure
figure_4A = cbe_data %>%
  flushOutData(., "C") %>%
  filter(!is.na(Dinucleotide)) %>%
  ggplot(aes(x = Position, y = Edit_norm, fill = Dinucleotide)) +
  ylab("Percent Editing") +
  xlab("Position in Protospacer") +
  geom_bar(stat = "summary", fun.y = "mean", color  = "black", alpha = 0.7) +
  stat_smooth(aes(outfit=cbe_fit<<-..y..), span = 0.5, color = "black") + # 0.4 for abe and ~0.43 for
cbe
  geom_point(alpha = 0.7) +
  scale_y_continuous(limits  = c(-0.01, 1.01), breaks = seq(0,1, 0.2), labels =
scales::percent_format()) +
  scale_x_continuous(breaks = seq(0, 20, 2)) +
  coord_cartesian(xlim =  c(1,20), clip = "on")  +
  scale_fill_manual(values = c("All" = "white", "AC" = "#4daf4a", "CC" = "#377eb8", "GC" = "grey",
"TC" = "#e41a1c")) +
  labs(fill = "Pre-dinucleotide") +
  theme_bw(base_size = 18) +
  theme(aspect.ratio = 1/2,
        panel.grid.major = element_blank(), panel.grid.minor = element_blank()) +
  facet_grid(cols = vars(Dinucleotide))

## make a dataframe for predicting edits
cbe_prediction = data.frame(predicted_editing = cbe_fit) %>%
  # filter(Dinucleotide != "All") %>%
  #mutate(Position = rep(1:80, times = 4)) %>%
  mutate(Position = rep(rep(1:20, each = 4), times = 5)) %>%
  mutate(letter = rep(rep(c("a", "b", "c", "d"), times = 20), times = 5)) %>%
  mutate(Dinucleotide = rep(c("All", "TC", "CC", "AC", "GC"), each = 80)) %>%
  filter(letter == "d") %>%
  dplyr::select(-letter) %>%
  mutate(predicted_editing = {ifelse(predicted_editing < 0, 0, predicted_editing)}) %>%
  mutate(Enzyme = "BE4")

### Figure 4B
### ABE base editing by position
```

```
## Load data
abe_data =
read_tsv("/Users/kluesner/Desktop/Research/spliceR/submissions/additional_files/abe_meta_data.tsv")
%>%

  mutate(Protospacer = toupper(Protospacer)) %>%
  mutate(Edit = `G`) %>%

  # Establish the maximum edit for each cell type in each paper
  group_by(`Cell type`, Paper) %>%
  mutate(paper_max = max(Edit)) %>%
  ungroup %>%

  # Normalize the edits to the max edit observed in the paper
  mutate(Edit_norm = Edit/paper_max) %>%

  # For each position in each unique guide, calculate the average normalized edit
  group_by(Position, Protospacer) %>%
  dplyr::summarise(Edit_norm = mean(Edit_norm)) %>%
  ungroup() %>%
  arrange(Protospacer)

## plot data

### Figure 4B
# ABE base editing by position, by dinucleotide
figure_4B = abe_data %>%
  flushOutData(., "A") %>%
  filter(!is.na(Dinucleotide)) %>%
  ggplot(aes(x = Position, y = Edit_norm, fill = Dinucleotide)) +
  ylab("Percent Editing") +
  xlab("Position in Protospacer") +
  geom_bar(stat = "summary", fun.y = "mean", color  = "black", alpha = 0.7) +
  stat_smooth(aes(outfit=abe_fit<<-..y..), span = 0.4, color = "black") + # 0.4 for abe and ~0.43 for
cbe
  geom_point() +
  scale_y_continuous(limits  = c(-0.01, 1.01), breaks = seq(0,1,0.2), labels =
scales::percent_format()) +
  scale_x_continuous(breaks = seq(0, 20, 2)) +
  coord_cartesian(xlim =  c(1,20), clip = "on")  +
  scale_fill_manual(values = c("All" = "white", "AA" = "#4daf4a", "CA" = "#377eb8", "GA" = "grey",
"TA" = "#e41a1c")) +
  labs(fill = "Pre-dinucleotide") +
  theme_bw(base_size = 18) +
  theme(aspect.ratio = 1/2,
        panel.grid.major = element_blank(), panel.grid.minor = element_blank()) +
  facet_grid(cols = vars(Dinucleotide))

## make a dataframe for predicting edits
abe_prediction = data.frame(predicted_editing = abe_fit) %>%
  # filter(Dinucleotide != "All") %>%
  #mutate(Position = rep(1:80, times = 4)) %>%
  mutate(Position = rep(rep(1:20, each = 4), times = 5)) %>%
  mutate(letter = rep(rep(c("a", "b", "c", "d"), times = 20), times = 5)) %>%
  mutate(Dinucleotide = rep(c("All", "TA", "CA", "GA", "AA"), each = 80)) %>%
  filter(letter == "d") %>%
  dplyr::select(-letter) %>%
  mutate(predicted_editing = {ifelse(predicted_editing < 0, 0, predicted_editing)}) %>%
  mutate(Enzyme = "ABE7.10")

### Figure 4C
# Logo plots of each guide

plotLogo = function(data, facet, method = "bit"){

  plotting_data = data %>%
    mutate(TargetMotif = toupper(substr(Protospacer, start = Position - 2, stop = Position + 2))) %>%
    mutate(TargetMotif = {ifelse(nchar(TargetMotif) < 5, paste0(TargetMotif, " "), TargetMotif)}) %>%
    inner_join(., data.frame(stringsAsFactors = F,
                         Motif = c("SA", "SD", "pmSTOP"), Motif_Name = c("Acceptor", "Donor",
```

```
"pmSTOP"))) %>%
    mutate(Facet = paste0(Enzyme, " ", Motif_Name))

  facets = unique(plotting_data$Facet)

  plotting_data %>%
    filter(Facet == facets[facet]) %>%
    filter(!is.na(TargetMotif) & nchar(TargetMotif) == 5) %>%
    dplyr::select(TargetMotif) %>%
    ggseqlogo::ggseqlogo(., method = method) +
    scale_y_continuous(limits = c(0,2), breaks = seq(0,2,0.5), labels = seq(0,2,0.5)) +
    scale_x_continuous(breaks = 1:5, labels = c("N-2", "N-1", "Target", "N+1", "N+2")) +
    ggtitle(facets[facet]) +
    # annotate(geom = "text", label = facets[facet], x = 0.5, y = 1.8, size = 8, hjust = 0) +
    theme_bw(base_size = 32) +
    theme(panel.grid = element_blank(), aspect.ratio = 1)
}

figure_4Ci = plotLogo(experimental_data, 1)
figure_4Cii = plotLogo(experimental_data, 2)
figure_4Ciii = plotLogo(experimental_data, 3)
figure_4Civ = plotLogo(experimental_data, 4)
figure_4Cv = plotLogo(experimental_data, 5)

### Figure 4 stats
n_guides = bind_rows(abe_data, cbe_data) %>%
  .$Protospacer %>%
  unique() %>%
  length

n_papers = bind_rows(abe_data, cbe_data) %>%
  .$Paper %>%
  unique() %>%
  length %>%
  {. + 1}

n_edits = bind_rows(abe_data, cbe_data) %>%
  dplyr::select(Protospacer, Position) %>%
  distinct %>%
  nrow()


##### FIGURE 5 ######
### Figure 5A
# Position dependent effets of correlation of base editing

position_cor_data = experimental_data %>%
  group_by(Exon_Class) %>%
  dplyr::summarize(cor = signif(correlation(Edit, Protein_Loss), 3),
                   pvalue = signif(correlation_pvalue(Edit, Protein_Loss), 3)) %>%
  mutate(cor = paste0("r = ", cor)) %>%
  mutate(pvalue = paste0("p-value = ", pvalue))

figure_5A = experimental_data %>%
  filter(Experiment != "STE2") %>%
  # mutate(Enzyme_Motif = factor()) $%>
  ggplot(aes(x  = Edit, y = Protein_Loss, fill = Exon_Class)) +
  geom_abline(slope = 1, intercept = 0, linetype = "dashed") +
  geom_smooth(method = "lm", color = "black", aes(fill = NULL)) +
  geom_point(pch = 21) +
  scale_y_continuous(labels = scales::percent_format(), limits = c(0, 1)) +
  scale_x_continuous(labels = scales::percent_format(), limits = c(0, 1)) +
  theme_bw(base_size = 12) +
  geom_label(data = position_cor_data, aes(label = cor, fill = NULL), x = 0.25, y = 0.75) +
  xlab("Genetic Editing") +
  ylab("Protein Loss") +
  facet_grid(cols = vars(Exon_Class)) +
  theme(aspect.ratio = 1,
        axis.text.x = element_text(hjust = 1, angle = 45),
        legend.position = "none",
```

```
        panel.grid.major = element_blank(),
        panel.grid.minor = element_blank()
  )

### Figure 5B
figure_5B = experimental_data %>%
  filter(Experiment != "STE2") %>%
  mutate(diff = abs(Protein_Loss - Edit)) %>%
  ggplot(aes(x = Exon_Class, y = diff, fill = Exon_Class)) +
  geom_boxplot(outlier.alpha = 0, alpha = 0.7) +
  geom_line(aes(x = as.numeric(Exon_Class), y = diff, fill = NULL), fun.y = "median", stat =
"summary", lwd = 1.5) +
  geom_point() +
  ylab("Error (|Protein Loss - Editing|)") +
  xlab("Exon") +
  scale_y_continuous(limits = c(0,1), breaks = seq(0,1, 0.2), labels = scales::percent_format()) +
  theme_classic(base_size = 24) +
  theme(legend.position = "none")

exon_group.lm = experimental_data %>%
  filter(Experiment != "STE2") %>%
  mutate(diff = abs(Protein_Loss - Edit)) %>%
  lm(diff ~ 0 + Exon_Group, data = .)

table_5B = exon_group.lm %>%
  summary() %>%
  .[[4]] %>%
  as.data.frame() %>%
  rownames_to_column("Exon_Group") %>%
  mutate(Exon_Group = gsub("Exon_Group", "", Exon_Group)) %>%
  as_tibble

### Figure 5C
domain_cor_data = experimental_data %>%
  filter(Experiment != "STE2") %>%
  filter(Domain != "UTR") %>%
  group_by(Domain) %>%
  dplyr::summarize(cor = signif(correlation(Edit, Protein_Loss), 3),
                   pvalue = signif(correlation_pvalue(Edit, Protein_Loss), 3)) %>%
  mutate(cor = paste0("r = ", cor)) %>%
  mutate(pvalue = paste0("p-value = ", pvalue))

figure_5C = experimental_data %>%
  filter(Experiment != "STE2") %>%
  filter(Domain != "UTR") %>%
  mutate(Domain = factor(Domain, levels = c("Extracellular", "Transmembrane", "Intracellular"))) %>%
  ggplot(aes(x  = Edit, y = Protein_Loss, fill = Domain)) +
  geom_abline(slope = 1, intercept = 0, linetype = "dashed") +
  geom_smooth(method = "lm", color = "black", aes(fill = NULL)) +
  geom_point(pch = 21, size = 2) +
  scale_y_continuous(labels = scales::percent_format(), limits = c(0, 1)) +
  scale_x_continuous(labels = scales::percent_format(), limits = c(0, 1)) +
  theme_bw(base_size = 18) +
  geom_label(data = domain_cor_data, aes(label = cor, fill = NULL), x = 0.25, y = 0.75) +
  xlab("Genetic Editing") +
  ylab("Protein Loss") +
  facet_grid(.~Domain) +
  theme(aspect.ratio = 1,
        axis.text.x = element_text(hjust = 1, angle = 45),
        legend.position = "none",
        panel.grid.major = element_blank(),
        panel.grid.minor = element_blank()
  )


##### Discussion #####
### Calculate breakdown of splice site mutations that are in the splice acceptor
clinvar =
read_tsv("/Users/kluesner/Desktop/Research/spliceR/submissions/additional_files/clinvar_20191223.vcf",
skip = 27)
```

```
splice_mutations = clinvar %>%
  mutate(splice = {ifelse(grepl("splice_donor", INFO), "donor",
                          ifelse(grepl("splice_acceptor", INFO), "acceptor","none"))}) %>%
  dplyr::select(CHROM, POS, ID, REF, ALT, splice) %>%
  dplyr::group_by(splice) %>%
  summarize(mutations = length(splice)) %>%
  ungroup() %>%
  mutate(total = sum(mutations), perc = scales::percent(mutations/total))



####### Supplementary Figures #######

##### FIGURE S1 #####

### Figure S1A
# Diagram of SpliceR algorithm

##### FIGURE S2 AND S3 #####
supplementaryFigure = function(enzyme, motif){
  data_tmp = guides
  # enzyme = "ABE"
  # motif = "acceptor"

  ## Establish temporary dataframe
  message("filtering data.")
  figure_data_tmp = data_tmp %>%
    # filter on desired enzyme
    filter(grepl(enzyme, be)) %>%
    # filter on desired splice_site
    filter(splice_site == motif) %>%
    # duplicate the data for NGN pam for aggregating and plotting purposes
    mutate(NGN = "NGN") %>%
    gather(type, PAM, c("pam_class", "NGN")) %>%
    group_by(gene_id, splice_site, PAM) %>%
    # summarize the earliest guide and number of guides for each gene for each pam
    dplyr::summarise(earliest_guide = min(position_score), number_of_guides = length(position_score))
%>%
    # factor for plotting order
    mutate(PAM = factor(PAM, levels = c("NGG", "NGA", "NGT", "NGC", "NGN")))

  # Change enzyme name for plotting
  enzyme = if(enzyme == "ABE") {"ABE"} else {"CBE"}

  ## Plotting
  # define the axis adjustment scalar
  axis_adjustment = sum(nsesi_gene_ids %in% (figure_data_tmp %>% .$gene_id))/length(nsesi_gene_ids)
  # define the earliest percent distance into the transcript 50% of the guides have
  guide50_tmp = figure_data_tmp %>% filter(PAM == "NGN") %>% .$earliest_guide %>% median
  # define the number of guides 50% of genes have
  number50_tmp = figure_data_tmp %>% filter(PAM == "NGN") %>% .$number_of_guides %>% median

  # Plot the cumulative earliest guide position
  message("making earliest guide plot")
  figure_data_tmp %>%
    ggplot(aes(x = earliest_guide, color = PAM)) +
    stat_ecdf(geom = "step", pad = FALSE, lwd = 1.5, alpha = 0.8) +
    scale_y_continuous(limits = c(0,1)/axis_adjustment, breaks = seq(0,1,0.2)/axis_adjustment, labels
= paste0(seq(0,1,0.2)*100, "%")) +
    scale_x_continuous(limits = c(0,1), breaks = seq(0, 1, 0.2)) +
    ggtitle(paste0(enzyme, " splice ", motif, " sgRNAs")) +
    ylab("Cumulative % of targetable genes") +
    xlab("Earliest BE-Splice targetable splice-site\n(relative position in mRNA)") +
    coord_cartesian(xlim = c(0, 1.0), ylim = c(0, 1)/axis_adjustment, expand = F) +
    theme_bw(base_size = 24) +
    theme(panel.border = element_rect(colour = "black", fill=NA),
          aspect.ratio = 1,
          legend.position = c(0.825,  0.25/axis_adjustment),
          legend.background = element_rect(fill = "white", color = "black"),
          plot.margin = unit(c(1,1,1,1), "cm")
```

```
    ) +
    labs(color = "PAM") +
    scale_color_manual(values = colors) +
    geom_hline(yintercept = axis_adjustment/axis_adjustment, linetype = "dashed") +
    annotate(geom = "text", size = 5, x = if(motif == "acceptor"){0.2}else{0.8}, y = 0.90, label =
paste0(as.percent(axis_adjustment)," of genes\nare targettable")) +
    annotate(geom = "text", size = 5, x = if(motif == "acceptor"){0.2}else{0.8}, y = 0.625, label =
paste0("50% of earliest guides\nare ", as.percent(guide50_tmp), " into the\ntranscript or earlier")) +
    annotate(geom = "segment",  x = guide50_tmp, xend = guide50_tmp, y = 0, yend = 0.5, linetype =
"dashed") +
    annotate(geom = "segment",  x = 0, xend = guide50_tmp, y = 0.5, yend = 0.5, linetype = "dashed")
  ggsave(paste0("/Users/kluesner/Desktop/Research/spliceR/submissions/figures/figureS2/", enzyme, "_",
motif, "_earliestGuide.tiff"), device = "tiff")

  figure_data_tmp %>%
    ggplot(aes(x = number_of_guides, color = PAM)) +
    stat_ecdf(geom = "step", pad = FALSE, lwd = 1.5, alpha = 0.9) +
    scale_y_continuous(limits = c(0,1)/axis_adjustment, breaks = seq(0,1,0.2)/axis_adjustment, labels
= paste0(seq(0,1,0.2)*100, "%")) +
    scale_x_log10() +
    annotation_logticks(sides = "b") +
    ylab("Cumulative % of targetable genes") +
    xlab("Number of BE-Splice gRNAs per gene") +
    ggtitle(paste0(enzyme, " splice ", motif, " sgRNAs")) +
    coord_cartesian(xlim = c(0.9999, max(figure_data_tmp$number_of_guides)), ylim = c(-0.005,
1.005/axis_adjustment), expand = F) +
    theme_bw(base_size = 24) +
    theme(panel.border = element_rect(colour = "black", fill=NA),
          aspect.ratio = 1,
          legend.position = c(0.8,  0.25/axis_adjustment),
          legend.background = element_rect(fill = "white", color = "black"),
          plot.margin = unit(c(1,1,1,1), "cm")
    ) +
    labs(color = "PAM") +
    scale_color_manual(values = colors) +
    geom_hline(yintercept = axis_adjustment/axis_adjustment, linetype = "dashed") +
    annotate(geom = "text", size = 5, x = 10^(0.8*log10(max(figure_data_tmp$number_of_guides))), y =
0.90, label = paste0(as.percent(axis_adjustment)," of genes\nare targettable")) +
    annotate(geom = "text", size = 5, x = 10^(0.8*log10(max(figure_data_tmp$number_of_guides))), y =
0.625, label = paste0("50% of targeted genes\nhave ", number50_tmp, " guides or more\ntargeting
them")) +
    annotate(geom = "segment",  x = number50_tmp, xend = number50_tmp, y = 0, yend = 0.5, linetype =
"dashed") +
    annotate(geom = "segment",  x = 0, xend = number50_tmp, y = 0.5, yend = 0.5, linetype = "dashed")
  ggsave(paste0("/Users/kluesner/Desktop/Research/spliceR/submissions/figures/figureS3/", enzyme, "_",
motif, "_numberOfGuides.tiff"), device = "tiff")
}

supplementaryFigure("BE3", "donor")
supplementaryFigure("BE3", "acceptor")
supplementaryFigure("ABE", "donor")
supplementaryFigure("ABE", "acceptor")

### Figure S2A
# CBE splice donor sgRNAs: cumsum vs. n per gene

### Figure S2B
# ABE splice donor sgRNAs: cumsum vs. n per gene

### Figure S2C
# CBE splice donor sgRNAs: cumsum vs. n per gene

### Figure S2D
# ABE splice donor sgRNAs: cumsum vs. n per gene

### Figure S3A
# CBE splice donor sgRNAs: cumsum vs. earliest guide

### Figure S3B
# ABE splice donor sgRNAs: cumsum vs. earliest guide
```

```
### Figure S3C
# CBE splice donor sgRNAs: cumsum vs. earliest guide

### Figure S3D
# ABE splice donor sgRNAs: cumsum vs. earliest guide

##### FIGURE S4 #####

### Figure S4A
# Representative gating strategy

##### FIGURE S5 #####

### Figure S5A
# CBE: Same analysis of position dependent context, but with only the post dinucleotide
figure_S5C = cbe_data %>%
  flushOutData(., "C") %>%
  filter(!is.na(PostDinucleotide)) %>%
  ggplot(aes(x = Position, y = Edit_norm, fill = PostDinucleotide)) +
  ylab("Percent Editing") +
  xlab("Position in Protospacer") +
  geom_bar(stat = "summary", fun.y = "mean", color  = "black", alpha = 0.7) +
  stat_smooth(aes(outfit=tmp_fit<<-..y..), span = 0.5, color = "black") + # 0.4 for abe and ~0.43 for
cbe
  geom_point() +
  scale_y_continuous(limits  = c(0, 1), breaks = seq(0,1, 0.2), labels = scales::percent_format()) +
  scale_x_continuous(breaks = seq(0, 20, 2)) +
  coord_cartesian(xlim =  c(1,20), clip = "on")  +
  scale_fill_manual(values = c("All" = "white", "CA" = "#4daf4a", "CC" = "#377eb8", "CG" = "grey",
"CT" = "#e41a1c")) +
  labs(fill = "Post-dinucleotide") +
  theme_bw(base_size = 18) +
  theme(aspect.ratio = 1/2,
        panel.grid.major = element_blank(), panel.grid.minor = element_blank()) +
  facet_grid(cols = vars(PostDinucleotide))

### Tables
figure_S5A = cbe_data %>%
  flushOutData(., "C")  %>%
  lm(Edit_norm ~ 0 + Dinucleotide, data = .) %>%
  summary() %>%
  .[[4]] %>%
  as.data.frame() %>%
  mutate(Dinucleotide = c("All", "TC", "CC", "AC", "GC")) %>%
  dplyr::rename(`Average Editing` = Estimate) %>%
  dplyr::select(Dinucleotide, everything()) %>%
  mutate(`Average Editing` = scales::percent(`Average Editing`),
         `Std. Error` = scales::percent(`Std. Error`),
         `t value` = signif(`t value`, 3),
         `Pr(>|t|)` = signif(`Pr(>|t|)`, 3)
  )

figure_S5E = cbe_data %>%
  flushOutData(., "C")  %>%
  lm(Edit_norm ~ 0 + PostDinucleotide, data = .) %>%
  summary() %>%
  .[[4]] %>%
  as.data.frame() %>%
  mutate(PostDinucleotide = c("All", "CT", "CC", "CA", "CG")) %>%
  dplyr::rename(`Average Editing` = Estimate) %>%
  dplyr::select(PostDinucleotide, everything()) %>%
  mutate(`Average Editing` = scales::percent(`Average Editing`),
         `Std. Error` = scales::percent(`Std. Error`),
         `t value` = signif(`t value`, 3),
         `Pr(>|t|)` = signif(`Pr(>|t|)`, 3)
  )


### Figure S5B
```

```
# ABE: Same analysis of position dependent context, but with only the post dinucleotide
figure_S5D = abe_data %>%
  flushOutData(., "A") %>%
  filter(!is.na(PostDinucleotide)) %>%
  ggplot(aes(x = Position, y = Edit_norm, fill = PostDinucleotide)) +
  ylab("Percent Editing") +
  xlab("Position in Protospacer") +
  geom_bar(stat = "summary", fun.y = "mean", color  = "black", alpha = 0.7) +
  stat_smooth(aes(outfit=tmp_fit<<-..y..), span = 0.4, color = "black") + # 0.4 for abe and ~0.43 for
cbe
  geom_point() +
  scale_y_continuous(limits  = c(0, 1), breaks = seq(0,1, 0.2), labels = scales::percent_format()) +
  scale_x_continuous(breaks = seq(0, 20, 2)) +
  coord_cartesian(xlim =  c(1,20), clip = "on")  +
  scale_fill_manual(values = c("All" = "white", "AA" = "#4daf4a", "AC" = "#377eb8", "AG" = "grey",
"AT" = "#e41a1c")) +
  labs(fill = "Post-dinucleotide") +
  theme_bw(base_size = 18) +
  theme(aspect.ratio = 1/2,
        panel.grid.major = element_blank(), panel.grid.minor = element_blank()) +
  facet_grid(cols = vars(PostDinucleotide))


### tables
figure_S5B = abe_data %>%
  flushOutData(., "A")  %>%
  lm(Edit_norm ~ 0 + Dinucleotide, data = .) %>%
  summary() %>%
  .[[4]] %>%
  as.data.frame() %>%
  mutate(Dinucleotide = c("All", "TA", "CA", "AA", "GA")) %>%
  dplyr::rename(`Average Editing` = Estimate) %>%
  dplyr::select(Dinucleotide, everything()) %>%
  mutate(`Average Editing` = scales::percent(`Average Editing`),
         `Std. Error` = scales::percent(`Std. Error`),
         `t value` = signif(`t value`, 3),
         `Pr(>|t|)` = signif(`Pr(>|t|)`, 3)
  )

figure_S5F =abe_data %>%
  flushOutData(., "A")  %>%
  lm(Edit_norm ~ 0 + PostDinucleotide, data = .) %>%
  summary() %>%
  .[[4]] %>%
  as.data.frame() %>%
  mutate(PostDinucleotide = c("All", "AT", "AC", "AA", "AG")) %>%
  dplyr::rename(`Average Editing` = Estimate) %>%
  dplyr::select(PostDinucleotide, everything()) %>%
  mutate(`Average Editing` = scales::percent(`Average Editing`),
         `Std. Error` = scales::percent(`Std. Error`),
         `t value` = signif(`t value`, 3),
         `Pr(>|t|)` = signif(`Pr(>|t|)`, 3)
  )


##### FIGURE S6 #####
# Gene maps with guides

##### FIGURE S7 - S13 #####
# B2M mapped
# TRAC mapped
# TRBC mapped
# CD3D mapped
# CD3E mapped
# CD3G mapped
# CD427 (CD3ζ) mapped

####### Saving Figures #######

##### FIGURE 1 #####
```

```
figure_1C %>%
  kableExtra::kable(., "latex", booktabs = T) %>%
  kableExtra::kable_styling(latex_options = c("striped", "scale_down")) %>%
  kableExtra::as_image(., file =
"/Users/kluesner/Desktop/Research/spliceR/submissions/figures/figure1/figure_1C.png")

ggsave("/Users/kluesner/Desktop/Research/spliceR/submissions/figures/figure1/figure_1D.tiff",
figure_1D)
ggsave("/Users/kluesner/Desktop/Research/spliceR/submissions/figures/figure1/figure_1E.tiff",
figure_1E)


##### Figure 2 #####
ggsave("/Users/kluesner/Desktop/Research/spliceR/submissions/figures/figure2/figure_2C.tiff",
figure_2C)

##### Figure 3 #####
ggsave("/Users/kluesner/Desktop/Research/spliceR/submissions/figures/figure3/figure_3A.tiff",
figure_3A, width = 22, height = 10, units = "in")
ggsave("/Users/kluesner/Desktop/Research/spliceR/submissions/figures/figure3/figure_3B.tiff",
figure_3B)
ggsave("/Users/kluesner/Desktop/Research/spliceR/submissions/figures/figure3/figure_3C.tiff",
figure_3C)
ggsave("/Users/kluesner/Desktop/Research/spliceR/submissions/figures/figure3/figure_3D.tiff",
figure_3D)


##### Figure 4 #####
ggsave("/Users/kluesner/Desktop/Research/spliceR/submissions/figures/figure4/figure_4A.tiff",
figure_4A, height = 12, width  = 20)
ggsave("/Users/kluesner/Desktop/Research/spliceR/submissions/figures/figure4/figure_4B.tiff",
figure_4B, height = 12, width  = 20)
ggsave("/Users/kluesner/Desktop/Research/spliceR/submissions/figures/figure4/figure_4Ci.tiff",
figure_4Ci)
ggsave("/Users/kluesner/Desktop/Research/spliceR/submissions/figures/figure4/figure_4Cii.tiff",
figure_4Cii)
ggsave("/Users/kluesner/Desktop/Research/spliceR/submissions/figures/figure4/figure_4Ciii.tiff",
figure_4Ciii)
ggsave("/Users/kluesner/Desktop/Research/spliceR/submissions/figures/figure4/figure_4Civ.tiff",
figure_4Civ)
ggsave("/Users/kluesner/Desktop/Research/spliceR/submissions/figures/figure4/figure_4Cv.tiff",
figure_4Cv)

##### Figure 5 #####
ggsave("/Users/kluesner/Desktop/Research/spliceR/submissions/figures/figure5/figure_5A.tiff",
figure_5A)
ggsave("/Users/kluesner/Desktop/Research/spliceR/submissions/figures/figure5/figure_5B.tiff",
figure_5B, height = 8)
ggsave("/Users/kluesner/Desktop/Research/spliceR/submissions/figures/figure5/figure_5C.tiff",
figure_5C)

##### Figure S5 #####
figure_S5A %>%
  kableExtra::kable(., "latex", booktabs = T) %>%
  kableExtra::kable_styling(latex_options = c("striped", "scale_down")) %>%
  kableExtra::as_image(., file =
"/Users/kluesner/Desktop/Research/spliceR/submissions/figures/figureS5/figure_S5A.png")

figure_S5B %>%
  kableExtra::kable(., "latex", booktabs = T) %>%
  kableExtra::kable_styling(latex_options = c("striped", "scale_down")) %>%
  kableExtra::as_image(., file =
"/Users/kluesner/Desktop/Research/spliceR/submissions/figures/figureS5/figure_S5B.png")

ggsave("/Users/kluesner/Desktop/Research/spliceR/submissions/figures/figureS5/figure_S5C.tiff",
figure_S5C, height = 12, width  = 20)
ggsave("/Users/kluesner/Desktop/Research/spliceR/submissions/figures/figureS5/figure_S5D.tiff",
figure_S5D, height = 12, width  = 20)

figure_S5E %>%
```

```
  kableExtra::kable(., "latex", booktabs = T) %>%
  kableExtra::kable_styling(latex_options = c("striped", "scale_down")) %>%
  kableExtra::as_image(., file =
"/Users/kluesner/Desktop/Research/spliceR/submissions/figures/figureS5/figure_S5E.png")

figure_S5F %>%
  kableExtra::kable(., "latex", booktabs = T) %>%
  kableExtra::kable_styling(latex_options = c("striped", "scale_down")) %>%
  kableExtra::as_image(., file =
"/Users/kluesner/Desktop/Research/spliceR/submissions/figures/figureS5/figure_S5F.png")
```
